# Evolution of Tunicate lifestyles shaped by *Myosin heavy chain* gene duplications, losses, and the diversification of tail muscle cell identities in *Oikopleura dioica*

**DOI:** 10.64898/2025.12.09.693291

**Authors:** Marc Fabregà-Torrus, Alfonso Ferrández-Roldán, Gaspar Sánchez-Serna, Beatriz Iralde Cárdenas, Cristian Cañestro

## Abstract

Tunicates offer a suitable system to study the evolutionary mechanisms underlying lifestyle transitions, ranging from the biphasic lifestyle of ascidians alternating swimming larva with sessile adults, to the fully free-swimming appendicularians. The appendicularian *Oikopleura dioica*, despite having only ten pairs of tail muscle cells, roughly half than in ascidian larvae, exhibits a remarkably rich repertoire of behaviours required for house inflation, swimming, nodding, and the rhythmic water pumping through the house. This apparent paradox suggests that functional diversification might have arose not from increased cell number, but from molecular specialization within a minimal musculature. To address this question, here, we reconstruct the evolution of the *Myosin class II heavy-chain* (*Myh*) gene family across tunicates and generated a developmental expression atlas of all *O. dioica Myh* genes. Phylogenetic analysis reveals that the cardio-paraxial *Myh* subfamily duplicated into two tunicate-specific subfamilies, followed by independent bursts of paralogue duplications in ascidians and appendicularians. In *O. dioica*, two distinct *Myh-Scp-Tb* paralogs show cardiac expression, and surprisingly, combinations of different *Myh-Scp-Tb* paralogues define multiple muscle cell identities along the tail. The innovation of this anteroposterior muscle regionalization associated to the *Myh-Scp-Tb* expansion provides a plausible mechanism for fine-scale modulation of contractile properties within a minimal musculature, helping to resolve the paradox of “fewer cells, but more contractile properties” in appendicularian tail movements. Conversely, the loss of the *Myh-Sj/bw* gene, which in ascidians is expressed with post-metamorphic body-wall and siphon muscles, is consistent with a pattern of regressive evolution associated to the absence of trunk muscles in *O. dioica*. This finding supports the view that the appendicularian lifestyle is a secondary derived condition, and that the last common ancestor of tunicates likely possessed an ascidian-like biphasic lifestyle. Together, our findings indicate that both gene-family duplication and gene loss have shaped the evolution of appendicularian tail muscle, enabling the emergence of complex tail-driven behaviours, and offering new insights into the ancestral lifestyle of tunicates.

## INTRODUCTION

Understanding how evolutionary transitions arise remains a central challenge in evolutionary biology, as these transitions represent pivotal points that increase biodiversity through novel ecological adaptations. Classical “major transitions” sensu Maynard Smith & Szathmáry, such as the emergence of eukaryotic cells or multicellularity, illustrate how profound reorganizations of biological systems can give rise to novel levels of biological complexity (Szathmáry, 2015). Beyond these canonical examples, numerous lineage-specific “great transitions” have also driven macroevolutionary shifts by enabling organisms to exploit new ecological opportunities through changes in lifestyle strategies.

Lifestyle transitions often involve extensive modifications of anatomy, physiology, and behaviour. In metazoans, these shifts are frequently mediated by metamorphosis, a developmental process that generates distinct larval and adult forms adapted to different ecological conditions (Fuchs et al., 2014; Laudet, 2011). Understanding the changes of the developmental and genetic mechanisms enabling such reorganizations is therefore essential for explaining how lifestyle transitions evolve.

In this context, tunicates, the sister group of vertebrates, comprising by ascidians, thaliaceans and appendicularians, stand out as a suitable system in which to study the evolutionary mechanisms underlying lifestyle transitions. Ascidians display a biphasic lifestyle, with a free-swimming, chordate-like larva that undergoes a dramatic metamorphosis to become a sessile adult (Lemaire, 2011; Satoh, 1994). Thaliaceans, which phylogenomics analyses robustly place within ascidians (Delsuc et al., 2018; Kocot et al., 2018), provide a paradigmatic example of a great transition from a biphasic, ascidian-like ancestor to a fully pelagic lifestyle, evolving jet-propelled locomotion, coloniality, and alternation of sexual and asexual generations (Bone, 1998). On the other hand, appendicularians occupy a strategic phylogenetic position for addressing the long-standing question of how was the lifestyle of the last common ancestor of tunicates, which is crucial to infer how was the olfactore ancestor from which vertebrates evolved. Although most phylogenomic studies consistently recover appendicularians as the sister group to all other tunicates (Delsuc et al., 2018; Kocot et al., 2018; Swalla et al., 2000; Wada, 1998), their extremely long branches and accelerated rates of molecular evolution have raised the possibility of artefactual basal position in the tunicate tree due to long-branch attraction (Singh et al., 2009; Stach, 2002; Yokobori et al., 2006). This ambiguity has fuelled a major debate over whether the last common ancestor of tunicates was free-swimming and appendicularian-like or biphasic with a sessile ascidian-like adult. Two contrasting hypotheses have emerged. *(1)* The free-swimming appendicularian-like ancestor hypothesis proposes that a completely free-swimming, tadpole-like organism represents the ancestral tunicate condition. This view is supported by parsimony, by the basal branching position of appendicularians in most phylogenomic trees, and by the fact that both cephalochordates and vertebrates are also fully motile chordates (Bourlat et al., 2006; Delsuc et al., 2006; Swalla et al., 2000; Wada, 1998). Under this scenario, sessility evolved only once, after the divergence of appendicularians in the lineage leading to ascidians + thaliaceans, with thaliaceans later returning secondarily to the pelagic condition. *(2)* The biphasic ascidian-like ancestor hypothesis, however, cannot be discarded. This alternative view, although less parsimonious, suggests that the tunicate ancestor already possessed an ascidian-like biphasic life cycle, with a swimming larva and a sessile adult, and that appendicularians subsequently evolved by neoteny, losing the drastic metamorphosis and adult sessile phase (Razy-Krajka & Stolfi, 2019; Stach et al., 2008). The hypothesis has gained recent support from several independent lines of evidence, including the extensive gene losses and developmental simplifications of appendicularians, which are more consistent with a derived, not ancestral, condition (Albalat & Cañestro, 2016). Notably, Ferrández-Roldán et al., (2021) showed that appendicularian *Oikopleura dioica* has lost many core components of a "deconstructed" cardiopharyngeal gene regulatory network, including key cardiac and pharyngeal muscle regulators, resulting in a fundamentally simplified cardiopharyngeal system with no siphon muscles and a simple laminar heart that beats against the stomach. The massive gene losses underlying the GRN deconstruction align with the absence of several ascidian features in appendicularians, strongly suggesting that the last common ancestor of tunicates likely had an ascidian-like biphasic lifestyle.

Among the features that enabled the emergence of the fully free-swimming appendicularian lifestyle, two evolutionary innovations stand out: the house and the tail maintenance throughout the life cycle. The “house” is a sophisticated filter-feeding mucous structure that appendicularians inflate, inhabit, abandon, and rebuild repeatedly throughout their life cycle (Fenaux, 1986; Flood & Deibel, 1998; Mikhaleva et al., 2018). This unique extracellular apparatus not only permits efficient suspension feeding in the pelagic environment but also imposes specific mechanical demands, as house inflation, water pumping, and house-entry and -exit behaviours all rely on controlled tail movements. The second key innovation is the lifelong retention of a functional tail. Unlike ascidians, whose larval tail is reabsorbed during metamorphosis, appendicularians bypass this transformation, preserving a chordate-like axial tail that remains the principal locomotor organ throughout life. This tail generates a rich repertoire of stereotyped and ecologically essential movements, including house inflation, house-entry and -exit, swimming, nodding, rapid escape responses, and the rhythmic pumping behaviours required for house function (Ferrández-Roldán et al., 2019; Kreneisz & Glover, 2015).

Despite the behavioural sophistication of appendicularians, the tail musculature of *O. dioica* is remarkably simple. Each tail contains only ten pairs of muscle cells, arranged in two longitudinal bands flanking the notochord (Fenaux, 1998). This minimal architecture stands in stark contrast with ascidians, whose larval tails include more numerous muscle cells (e.g. 36 in *Ciona robusta*; 42 in *Halocynthia roretzi*) specialised in a stereotyped oscillatory swimming pattern that enables short-range dispersal prior to metamorphosis (Razy-Krajka & Stolfi, 2019). The paradox is that *O. dioica*, with far fewer muscle cells, produces a much richer repertoire of movements than ascidian larvae (Ferrández-Roldán et al., 2019; Kreneisz & Glover, 2015). This discrepancy suggests that appendicularians did not increase behavioural complexity through expanded musculature, but rather through the functional specialization of a highly streamlined set of muscle fibres.

In chordates, differences in contraction speed, force generation, and fatigue resistance among muscle fibres are largely determined by the repertoire of *Myosin class II heavy chain* (*Myh*) genes or isoforms they express. Myh proteins define the biochemical and kinetic properties of sarcomeric thick filaments, providing a molecular mechanism by which even morphologically simple muscles can acquire diverse functional capacities. The evolutionary potential of *Myh* gene diversification is well illustrated in vertebrates, in which repeated expansions of sarcomeric *Myh* families have generated subfamilies with multiple isoforms with distinct biochemical and contractile properties. These vertebrate paralogues underpin the specialization for instance of fast-twitch (MYH1/2/4), embryo development (MYH3), cardiac, (MYH6/7) and extraocular muscles (MYH13), enabling the emergence of complex and highly coordinated locomotor behaviours—including sustained swimming in fishes, rapid escape responses, powered flight in birds and bats, and fine motor control in tetrapods (reviewed in Schiaffino & Reggiani, 2011).The tight correlation between *Myh* expansions and the evolution of novel locomotor organizations highlights the capacity of this gene family to act as a substrate for functional innovation. Thus, to better understand the evolution of the lifestyle of the last common tunicate ancestor and the developmental adaptations that facilitated the evolution of fully free-swimming and complex repertoire of tail movements in appendicularians, here we focus on the evolution of the *Myh* family in tunicates and investigate the developmental expression atlas of the complete catalogue of *Myh* genes in *O. dioica*, as the appendicularian model. Our results reveal that the cardio-paraxial *Myh* genes, in parallel with vertebrates, have been duplicated into two tunicate subfamilies, which in turn have independently expanded and diversified into several paralogues in appendicularians and ascidians. Our analysis also reveals the loss of a *Myh* type that in ascidians is expressed in the body wall, altogether providing further support to the biphasic ascidian-like ancestor hypothesis.

## MATERIALS AND METHODS

### Laboratory culture of *Oikopleura dioica*

*O. dioica* specimens were obtained from laboratory-maintained colonies established over five years ago. The founder individuals were originally collected from the Mediterranean coast near Barcelona (Catalonia, Spain) using a plankton net approximately 200 meters offshore. Culturing procedures followed the protocol described in Martí-Solans et al., (2015).

This project did not raise ethical concerns, as experimentation involving aquatic invertebrates is not subject to animal experimentation regulations, in accordance with Real Decreto 53/2013 and Catalan Law 5/1995, DOGC2073,5172. Nevertheless, all experimental procedures complied with European Union (EU) guidelines for animal care and were formally approved by the Ethical Committee for Animal Experimentation of the University of Barcelona (CEEA-2009).

### Genome searches, gene identification and phylogenetic analysis

*Myh* genes in *O*. *dioica* were first identified in the reference database of *O. dioica* relative to the Norwegian population (http://oikoarrays.biology.uiowa.edu/Oiko/) (Danks et al., 2013; Denoeud et al., 2010) with BLASTp and tBLASTn, using as queries the Myh protein sequences from the vertebrate *Homo sapiens* and the tunicate *Ciona robusta*, as well as several other chordates. The corresponding orthologs were then identified in the telomere-to-telomere genomic assemblies of the Barcelona and Okinawa *O. dioica* species, and in the superscaffolded version of the Osaka *O. dioica* cryptic species using tBLASTn (Plessy et al., 2024); NCBI BioProject ID PRJEB55052, and assemblies in https://zenodo.org/records/10241527. In each population, the newly identified *Myh* genes were used to search for further paralogs using tBLASTn to obtain the final *Myh* catalogue (all loci identified are provided in **Supplementary Table 1**).

*Myh* genes in other appendicularian and ascidian species were identified using *O. dioica, Ciona robusta* and other chordate Myh proteins as queries for tBLASTn against publicly available genomes (i.e. *Oikopleura albicans* SCLG01000000, *Oikopleura vanhoeffeni* SCLH01000000) (Naville et al., 2019) and ANISEED (i.e. *Ciona robusta*, *Ciona intestinalis*, *Ciona savignyi*, *Phallusia mammillata*, *Halocynthia roretzi*, *Halocynthia aurantium*) (Brozovic et al., 2018). All loci identified are provided in **Supplementary Table 2 and 3**. In addition, the list of vertebrate and cephalochordate sequences used for phylogenetic analysis within chordates are provided in **Supplementary Table 4**.

Protein alignments were generated with MUSCLE and MAFFT implemented in Aliview v1.28 (Larsson, 2014), NGPhylogeny (Lemoine et al., 2019) and reviewed by hand. Phylogenetic trees were based on Maximum Likelihood (ML) inferences calculated with PhyML v3.0 (Guindon et al., 2010), as well as IQ-Tree (Nguyen et al., 2015). LG was inferred as the best-fit substitution model for the Myh data according to both Akaike and Bayesian information criteria, with a gamma with 4 categories, estimated proportion of invariable sites of 0.1674, and a shape alpha of 0.9597 (Kalyaanamoorthy et al., 2017). Tree node support was inferred by fast likelihood-based methods aLRT SH-like, aLRT chi2-based and aBayes, and by standard or ultrafast bootstraps (n=100) according to computational capacity. Phylogenetic trees were inferred from alignments including the complete protein or just the most conserved head domain protein alignments, and in both cases producing the same tree topology regarding the *Myh* homology of appendicularian genes.

### Cloning and expression analyses

*O. dioica Myh* genes were PCR-amplified from cDNA or gDNA extracted from individuals of the Barcelona population, following the procedures described in Martí-Solans et al., (2016). PCR products were cloned using the TOPO TA Cloning Kit (K4530-20, Invitrogen), and the resulting plasmids were digested with the appropriate restriction enzymes to generate antisense DIG-labeled riboprobes for whole-mount in situ hybridization (WMISH) (Bassham & Postlethwait, 2000; Cañestro et al., 2007; Martí-Solans et al., 2016).

Due to the high sequence similarity among certain *Myh* coding regions, the design of specific probes was particularly challenging. To minimize potential cross-hybridization, probes were carefully designed to target the most divergent regions of each transcript. The fact that each probe showed a distinct expression pattern suggested a high probe specificity and a minimal cross-reactivity. Details on primers designed for the cloning of Myh genes are provided in **Supplementary Table 5**.

## RESULTS

### Phylogenetic analyses of the *Myh* gene catalogue across *O. dioica* populations

Systematic RBBH genomic searches and sequence alignments allowed us to identify the complete catalogue of *Myh* genes across four distinct *Oikopleura dioica* populations from different locations: Norway (NOR), Barcelona (BAR), Osaka (OSA), and Okinawa (OKI) (Denoeud et al., 2010; Plessy et al., 2024) **(Supplementary Table 1)**.

Phylogenetic analysis resolved eleven well-supported Myh paralogy groups within *O. dioica* populations (**Figure 1**). The explanation of gene nomenclature will be detailed later according to their relationships with chordate genes. Eight paralogy groups contained one representative of each population. These one-to-one orthologies consistently reconstruct the phylogenetic relationships among *O. dioica* populations, with BAR and NOR forming a clade, OSA as the next sister group, and OKI basally branching off, supporting that OSA is closer to BAR/NOR than to OKI, consistent with the presence of the Kuroshio current that separates Okinawa Island (OKI) from mainland Japan (OSA) (Masunaga et al., 2022).

**Figure 1.**
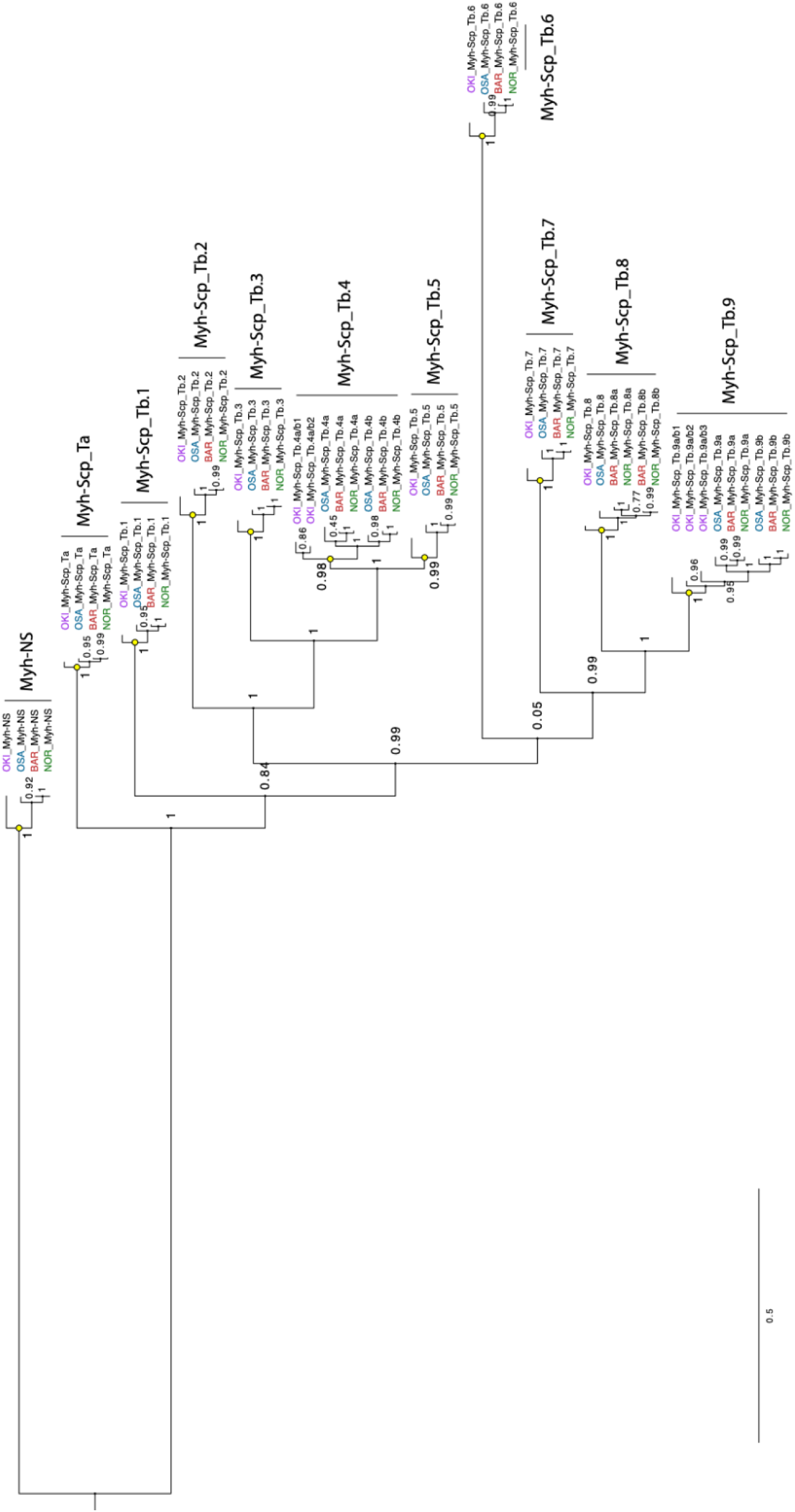
Maximum likelihood (ML) tree of all *O. dioica* class II *Myh* genes identified in four populations (BAR; NOR; OSA; OKI), based on full-length protein sequence alignment. Node support values indicate ultrafast bootstrap support. The scale bar indicates amino acid substitution.

However, three paralogy groups – *Myh-Scp_Tb.4*, *Myh-Scp_Tb.8*, and *Myh-Scp_Tb.9* – exhibited additional population-specific duplications or losses, indicating ongoing dynamic evolution of the *Myh* gene catalogue across *O. dioica* populations. In the *Myh-Scp_Tb.8* group, the phylogeny supported a basal duplication event in the BAR-NOR stem lineage (i.e. *Myh-Scp_Tb.8a* and *Myh-Scp_Tb.8b*) after divergence from the OSA lineage. In contrast, the *Myh-Scp_Tb.4* and *Myh-Scp_Tb.9* groups appeared to have undergone independent duplications events in the OKI lineage and in the common ancestor of OSA, BAR, and NOR. The short branch lengths observed in these paralogous groups suggested recent duplication events, although the possibility of ancestral duplications followed by subsequent gene loss cannot be excluded.

Comparative analyses of intron positions among *Myh* genes from all *O. dioica* populations revealed several distinctive intron-signatures unique to each of the eleven paralogy groups, providing strong support to the one-to-one orthologies as well as lineage-specific duplication or loss events (**Supplementary Figure 1**). The number of introns in *O. dioica Myh* genes was highly variable among paralogy groups, ranging from 2 to 14. Notably, the near-complete lack of conserved intron position among paralogy groups suggested that each group undergone extensive evolutionary divergence and did not support the possibility of gene conversion among *O. dioica Myh* paralogs.

These findings, revealing a remarkable genetic diversity in the *Myh* gene catalogue across *O. dioica* populations, reinforce the idea that the lineages represented by OKI, OSA, and BAR/NOR populations constitute, indeed, cryptic species, as it has been suggested by the massive genome scrambling observed among *O. dioica* populations (Plessy et al., 2024) and the lack of interbreeding between tested populations (Masunaga et al., 2022).

### Genome scrambling affects *Myh* macro- and micro-synteny conservation across populations

Macrosynteny analyses revealed a complete rearrangement of *Myh* loci throughout the genome among the three high quality genome assemblies at chromosome level (BAR, OSA, and OKI), with no evidence of the conserved *Myh* clustering patterns typically observed in other chordates (Weiss et al., 1999) (**Figure 2A**). This complete rearrangement limited the use of macrosynteny to support orthologies assignments, but instead it is consistent with the extreme genome scrambling described in *O. dioica* (Plessy et al., 2024), offering valuable insight of the effect of pervasive interchromosomal rearrangements, high rates of translocation, and erosion of large-scale syntenic blocks over gene families such as *Myh*.

**Figure 2.**
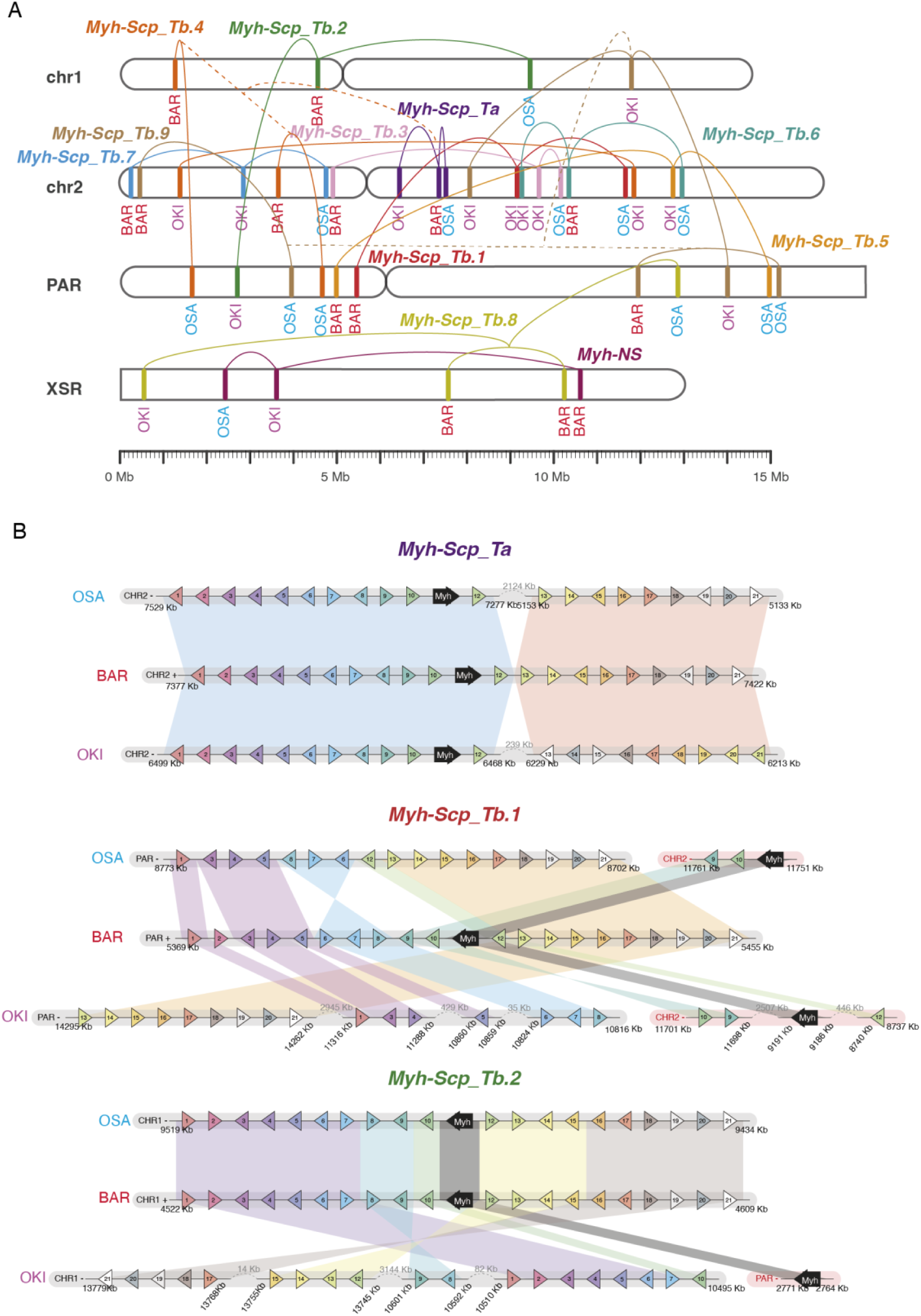
Comparative macrosynteny and microsynteny of the *Myh* catalogue among the three *O. dioica* cryptic species. (**A**) Comparative chromosome location of *Myh* genes with each paralog group labelled with a distinctive colour. (**B**) Three distinct examples of microsynteny conservation between the genomic neighbourhoods of *Myh* genes. The BAR genome was used as the reference in comparisons with OSA and OKI. The block containing the M*yh* genes (black arrow) and ten adjacent genes on each side were mapped in the OSA and OKI genomes.

For example, orthologs within the *Myh-Scp_Ta* group were located on the same chromosomal arm across all three populations suggesting a relatively conserved region on chromosome 2. In contrast, other orthologs were positioned on different chromosomal arms, including the examples of the orthologs of the *Myh-Scp_Tb.3* group, which were found on opposite chromosomal arms in BAR compared to OSA and OKI; or even the case of the *Myh-Scp_Tb.8*, which in OKI and BAR were located in chromosome XSR, but it had moved outside of the sex chromosome region in OSA, being found in the PAR region. Interestingly, our macrosyntenic analysis of the Myh also revealed several examples of interchromosomal gene translocations, events not observed in other analysed gene families (Plessy et al., 2024). Examples included the *Myh-Scp_Tb.1* group, in which the BAR ortholog was located on chromosome PAR, while its OSA and OKI counterparts were on chromosome 2; and the *Myh-Scp_Tb.2* group, where BAR and OSA orthologs were on chromosome 1, but the OKI ortholog was located on chromosome PAR.

Microsynteny analyses of orthologous subgroups provided further support to the orthologies shown in the phylogenetic tree (**Figure 2B**), while also revealing further details into the local structure of rearrangements. Microsynteny analyses showed cases of high conservation of gene synteny in the genomic neighbourhood of gene orthologs among BAR, OSA and OKI (i.e. *Myh-Scp_Ta*), cases of high conservation between BAR and OSA, but not OKI (i.e. *Myh-Scp_Tb.2*), and cases with low conservation among all populations (i.e. *Myh-Scp_Tb1*). In general, the degree of microsynteny conservation among *Myh* correlated with the phylogenetic distance between populations, once again supporting the positioning of OKI as the basal lineage. Interestingly, the fact that the level of synteny conservation was significantly variable across different Myh genomic neighbourhoods suggested that different evolutionary constraints may apply to each locus, as has been described for *Hox* and *Fgf* genes (Plessy et al., 2024).

### Chordate-wide phylogeny resolves major *Myh* lineage-specific duplications and losses across chordates

To investigate the evolutionary origin of the eight paralogy groups identified in *O. dioica*, we performed a tunicate-wide phylogenetic analysis including 2 additional appendicularian species and 6 ascidian species (**Supplementary Figure 2**). This analysis revealed the presence of 4 strongly supported *Myh* gene clusters in tunicates, according to the distribution of appendicularian and ascidian sequences throughout the tree. Three of these clusters included both ascidian and appendicularian genes, while the fourth cluster was composed exclusively of ascidian genes. This result suggested that the duplications that originated these clusters occurred were ancestral, likely before the divergence between ascidian and appendicularians.

To investigate the relationship of homology between those *Myh* tunicate clusters with the *Myh* families described in chordates, we performed a broad phylogenetic adding 76 Myh sequences representing the main vertebrate *Myh* subfamilies, all 27 cephalochordates *Myh* we found in the *Branchiostoma floridae* genome, and *Myh* sequences from BAR-*O. dioica* and *C. robusta* representing appendicularians and ascidians, respectively (**Figure 3**). Results revealed the presence of the two major *Myh* families in tunicates: *Non-sarcomeric Myh* (*Myh-NS*), and *Sarcomeric Myh* (*Myh-S*). The phylogenetic tree topology also revealed that *Myh-S* family had expanded by an ancestral duplication in the stem olfactore, before the vertebrate-tunicate split, giving rise to two major paralog types: the *jaw/body wall* type (*Myh-Sj/bw*) and the *cardioparaxial* type (*Myh-Scp*). The nomenclature adopted in this article for families, types and subfamilies of *Myh* follows the names of *Myh* classification in vertebrates (reviewed in Schiaffino & Reggiani, 2011) reconciled with our phylogenetic and expression patterns observed in non-vertebrate chordates.

**Figure 3.**
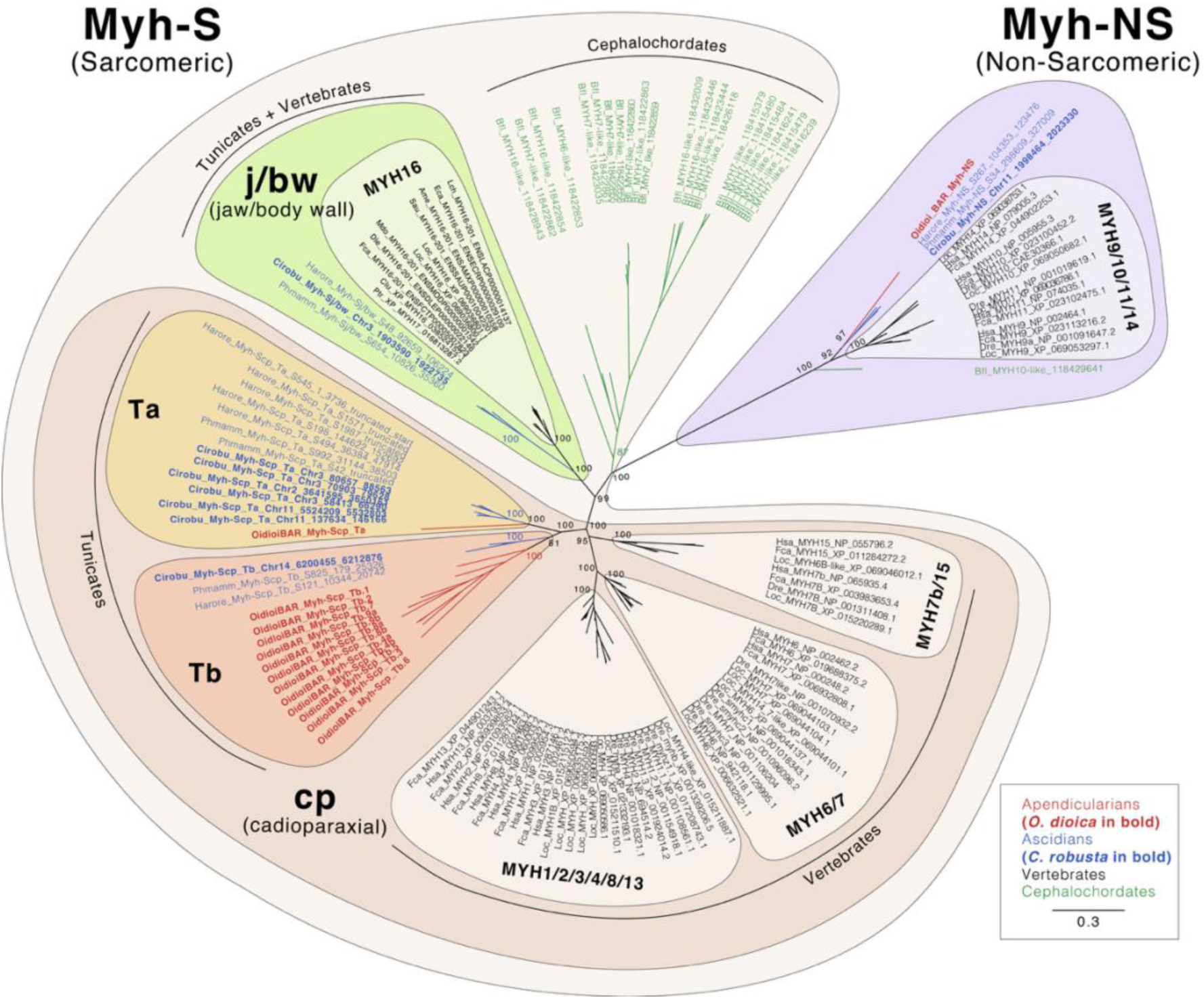
Evolutionary tree of *Myh* catalogue in chordates. ML phylogenetic tree of the *Myh* catalogue in chordates. Node support values indicate ultrafast bootstrap support. The scale bar indicates amino acid substitution. Species abbreviations: Vertebrates (in black): *Homo sapiens* (Has), Felis catus (Fca), Lepisosteus oculatus (Loc), Danio rerio (Dre), \**Canis lupus* (Clu), \**Pan troglodytes* (Ptr), \**Delphinapterus leucas* (Dle), \**Sparus aurata* (Sau), \**Astyanax mexicanus* (Ame), \**Erpetoichthys calabicurus* (Eca), \**Monodelphis domestica* (Mdo), \**Latimeria chalumnae* (Lch); Ascidian tunicates (in blue): *Ciona robusta* (Cirobu), *Ciona intestinalis* (Ciinte), *Ciona savignyi* (Cisavi), *Phallusia mammillata* (Phmamm), *Halocynthia roretzi* (Harore), *Halocynthia aurantium* (Haaura); Appendicularian tunicates (in red): *Oikopleura dioica* (Oidioi), *Oikopleura albicans* (Oialbi), *Oikopleura vanhoeffeni* (Oivanh); Cephalochordates (in green): *Branchiostoma floridae* (Bfl). For vertebrate species marked with an asterisk, only the *MYH16* group sequence was included.

#### · Non-Sarcomeric Myh gene family (Myh-NS)

*Myh-NS* gene family included paralogous *MYH9/10/11/14* subfamily of vertebrates, and only a single ortholog in all examined appendicularian and ascidian species, as well as one in cephalochordates. In vertebrates, this gene family is expressed in smooth muscle and non-muscle tissues, while in ascidians, the single ortholog it has been detected in the notochord and papillae precursors (Johnson et al., 2020; Nakayama-Ishimura et al., 2009)

#### · Sarcomeric jaw/body wall Myh type (Myh-Sj/bw)

The *Myh-Sj/bw* type included vertebrate *MYH16* genes and ascidian genes (Chiba et al., 2003), but no appendicularian sequences, suggesting therefore the loss of this *Myh* type in the appendicularian lineage after its divergence from ascidians. In vertebrates, *MYH16* is expressed in muscles associated to the jaw in the neck region, and its loss in the *Homo sapiens* primate ancestor is one of the most famous and paradigmatic example in which a gene loss could be adaptive by leading to the evolution of smaller jaw muscles and larger braincase that facilitated the evolution of human characteristics (Stedman et al., 2004). In ascidians, the *Myh-Sj/bw* type is expressed in post-metamorphic muscles of the body wall and siphons (Prünster et al., 2019) Therefore, the loss of this Myh type in appendicularians is consistent with the absence of body-wall and siphon muscles, and by extension, with the absence of a sessile, post-metamorphic adult stage.

#### · Sarcomeric cardioparaxial Myh type (Myh-Scp)

The phylogenetic tree revealed that the *Myh-Scp* type had been expanded by a tunicate-specific ancestral duplication into two paralogous subfamilies: the *Myh-Scp Tunicate_a* and *Tunicate_b* (*Myh-Scp_Ta* and *Myh-Scp_Tb*). This tunicate expansion was parallel to the expansions that gave rise to the Myh vertebrate subfamilies *MYH7b/15*, *MYH6/7* and *MYH1/2/3/4/8/13*, many of which had been associated with cardiac and paraxial muscle (Schiaffino & Reggiani, 2011). The *Myh-Scp_Ta* and *Myh-Scp_Tb* subfamilies were thus co-orthologs to all vertebrate *Myh-Scp* subfamilies.

Our phylogenetic analyses also revealed that the *Myh-Scp_Ta* and *Myh-Scp_Tb* subfamilies have suffered burst of duplications giving rise to multiple paralogs in each appendicularians and ascidian species. Interestingly, the burst of duplications seemed to follow opposite patterns of expansions in appendicularians and ascidians. All examined appendicularian species, for instance, contained multiple *Myh-Scp_Tb* genes, while only one *Myh-Scp_Tb* genes were present in most ascidians. In contrast, the *Myh-Scp_Ta* subfamily exhibited the opposite trend, in which ascidian species showed a substantial expansion of this subfamily (e.g. 7 in *C. intestinalis*, 6 in *C. robusta*, 7 in *C. savignyi*, 5 in *H. roretzi*, 5 in *H. aurantium*). In comparison, most appendicularian species contained fewer *Myh-Scp_Ta* (1 in *O. dioica* and 2 in *O. vanhoeffeni,* except for *O. albicans* that had 9). These results point that independent amplification of *Myh-Scp_Ta* and *Myh-Scp_Tb* subfamilies may account for multiple species-specific muscle adaptations in Tunicates, being *Myh-Scp_Tb* more prone to be expanded in appendicularians, and *Myh-Scp_Ta* ascidians.

The phylogenetic tree also showed important differences in branch lengths between the *Myh-Scp_Ta* and *Myh-Scp_Tb* subfamilies, being significantly longer in the latter, indicating lower amino acid sequence similarity among *Myh-Scp_Tb* paralogs. In contrast, many *Myh-Scp_Ta* paralogs in ascidians showed short branches and clustered together by species (e.g. *C. intestinalis* and *C. savignyi*), suggesting that many duplications were recent and species-specific. While the large differences in intron position among appendicularian *Myh* paralogs is not consistent with evolution by gene conversion, in the case of ascidians, their high sequence and structural similarity does not allow us to exclude the possibility of gene conversion in ascidian groups, a process described in the evolution of *Myh* genes in other lineages, contributing to the observed similarity (McGuigan, 2004). Comparative analyses of *Myh* intron positions revealed that while intron positions were vastly conserved between ascidians and vertebrates, almost none are conserved in *O. dioica* (**Supplementary Figure 1**). A few shared positions may reflect ancestral intron retention or recent parallel intron gains in conserved genomic hotspots. The intron position comparison also revealed a higher proportion of conserved introns within *Myh-NS* and *Myh-S groups*, providing additional support for the monophyly of these two major families, as well as for the close paralogous relationship between the *Myh-Sj/bw and the Myh-Scp*.

Taken together, all these contrasting patterns of duplication and sequence conservation among paralogs of the *Myh-Scp_Ta* and *Myh-Scp_Tb* subfamilies in ascidians and appendicularians likely suggested that distinct selective pressures may shape the evolution of each tunicate lineage, potentially linked to differences in the contractile demands of their respective lifestyles.

### Developmental expression atlas of the *Myh* gene catalogue in *O. dioica*

In ascidians, *Myh-Scp* gene expression had been associated to the heart and larval tail muscles (e.g. *Myh2* and *Myh1* in *B. schlosseri*; Prünster et al., (2019)). To better understand the lineage-specific expansions observed in appendicularians and to characterise the developmental deployment of the complete *O. dioica Myh* repertoire, we analysed the expression patterns of all 14 identified *Myh* genes in BAR population across embryonic and larval stages using whole-mount in situ hybridisation (WMISH). The resulting atlas revealed two major expression regimes that correspond to the two *Myh* families: one *Myh-NS* and the thirteen *Myh-Scp* paralogs, subdivided into one in the *Ta* and twelve in the *Tb* subfamilies.

#### · Non-muscle expression of Myh-NS single ortholog

The single *Myh-NS* gene of *O. dioica* displayed a broad non-muscle expression profile across development (**Figure 4 A**). Expression was first detected at the 100-cell stage in the nuclei of notochord precursor cells and persisted throughout tailbud stages, gradually diminishing during the hatchling stages as notochord vacuolisation progressed (**Figure 4 A1-A11**). No signal remained in the notochord by the pre-tailshift stage. At this later stage, *Myh-NS* expression became strongly upregulated in the tail epidermis and throughout the digestive system, including the mouth, pharyngeal epithelium, oesophagus, stomach lobes, and rectum (**Figure 4 A12**).

**Figure 4.**
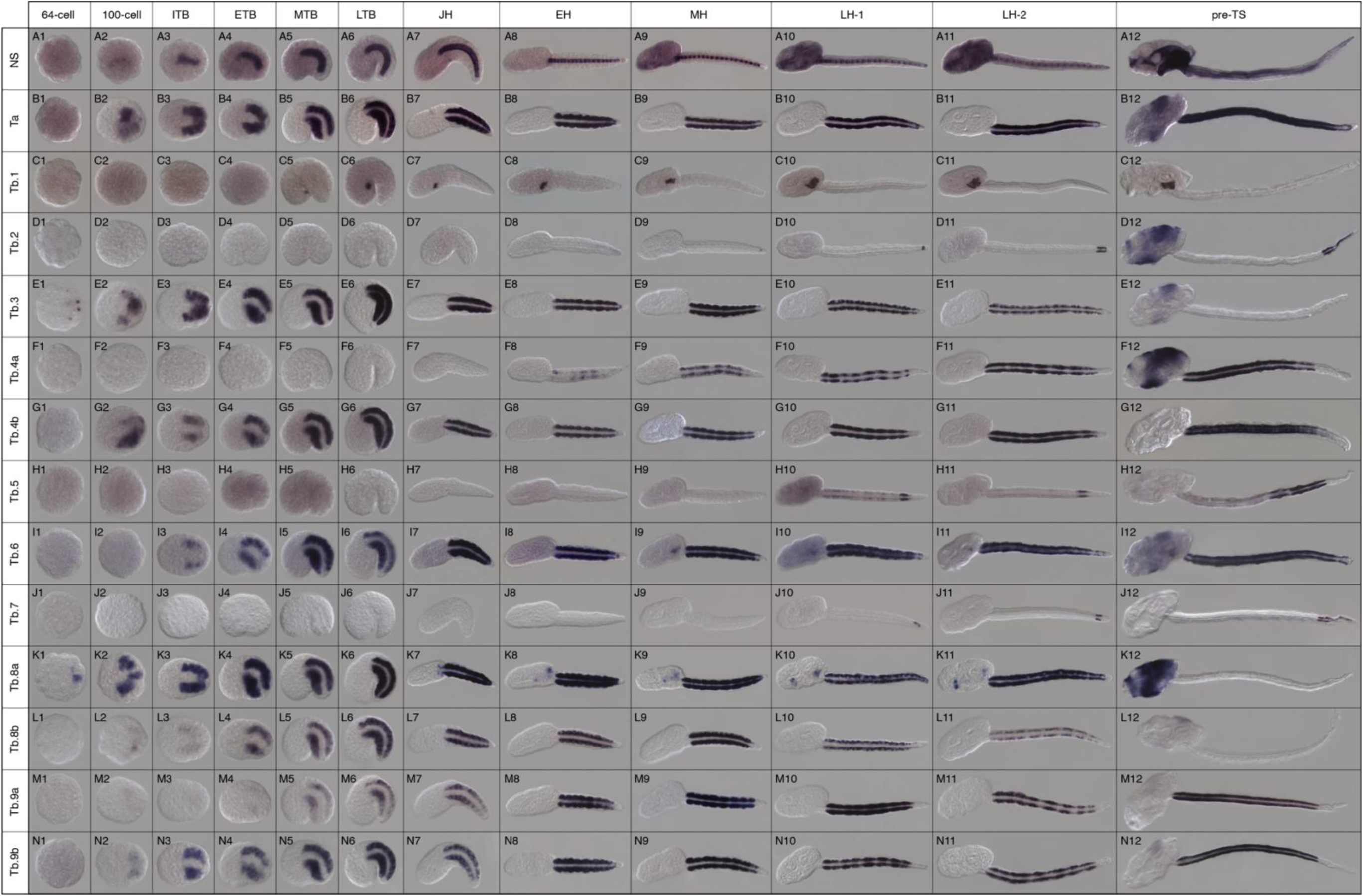
Developmental expression atlas of *O. dioica Myh* genes. Whole-mount in situ hybridization images of *O. dioica* at various developmental stages: 64-cell (A-N1), 100-cell (A-N2), incipient tailbud (ITB; A-N3), early tailbud (ETB; A-N4), mid tailbud (MTB; A-N5), late tailbud (LTB; A-N6), just hatchling (JH; A-N7), early hatchling (EH; A-N8), mid hatchling (MH; A-N9), late hatchling-1 (LH-1; A-N10), late hatchling-2 (LH-2; A-N11), pre-tailshift (pre-TS; A-N12). Images are left lateral views oriented anterior toward the left. *Myh-Scp_Ta* and *Myh-Scp_Tb*; referred to as *Ta* and *Tb* for the sake of simplicity.

#### · Paraxial muscle expression of the single ortholog for the *Myh-Scp_Ta* subfamily

The single *Myh-Scp_Ta* gene in *O. dioica* was exclusively expressed in all tail muscle cells throughout development, but not expressed in the heart (**Figure 4 B1-12**). Expression was first detected at 100-cell stage, prior to the early tailbud stage where eight muscles cell pairs were clearly identified (m1-8, **Figure 4 B2-B6**). After hatching, expression extended to the two most posterior muscle pairs, m9 and m10 (**Figure 4 B7-B12**). This pattern, involving all ten muscle cell pairs, persisted throughout the hatchling stages, including the pre-tailshift stage.

In ascidians, the *Myh-Scp_Ta* gene subfamily has suffered multiple independent expansions. Although a comprehensive analysis of expression across all paralogs is still lacking, Ta-type genes have been consistently reported to exhibit larval tail muscle-specific expression in various ascidian species (Chiba et al., 2003; Prünster et al., 2019; Razy-Krajka & Stolfi, 2019), which would be consistent with the expression of *Myh-Scp_Ta* in *O. dioica*.

#### · Temporal and spatial cardioparaxial expression regionalization of expanded paralogs for the *Myh-Scp_Tb* subfamily

The analysis of the 12 *Myh-Scp_Tb* paralogs of *O. dioica* revealed conserved cardioparaxial expression, although with unique and diversified expression patterns for each gene (**Figure 4 C1-N12**). These patterns included temporal differences (in the onset or downregulation of the expression at different developmental stages) and spatial differences (with expression restricted to distinct subsets of muscle cells along the anteroposterior axis).

##### Cardiac muscle expression

Two *Tb* paralogs, *Tb.1* and *Tb.6*, showed clear expression in heart muscle cells (**Figure 4 C, I1-12**). *Tb.1* was exclusively expressed in the heart, becoming the first robust heart-specific marker described in *O. dioica* (**Figure 4 C1-12**). Expression onset of *Tb.1* appeared when the precursor cardiac cells had completed their migration to the left ventral side of the trunk. *Tb.1* expression in myocardial cell was maintained throughout all hatchling stages, including the pre-tailshift stage, when the heart was already beating. In contrast to *Tb.1*, *Tb.6* was expressed in both cardiac and paraxial muscle (**Figure 4 I1-12**). *Tb.6* expression was initially restricted to paraxial muscle cells during tailbud stages, but later at the mid hatchling stage, *Tb.6* was also expressed in the heart (**Figure 4 I1-12**). The *Tb.6* signal in paraxial muscle cells included all cell pairs from m1 to m10, and the signal was consistently stronger in tail muscle cells than in heart cells.

##### Paraxial muscle expression

All *Tb* paralogs other than *Tb.1* and *Tb.6* were expressed in tail muscle cells and could be classified in two groups of expression: the “pan-muscle” group that were expressed in all 10 muscle pairs, and the “position-specific muscle” group that were expressed only in a subset of tail muscle cells. The pan-muscle group included *Tb.3* and *Tb.8a*, together with the already mentioned *Tb.6* (**Figure 4 E, I, K1-12**). The position-specific muscle included *Tb.4a*, *Tb.4b*, *Tb.8b*, *Tb.9a*, and *Tb.9b* that were expressed only in the first eight muscle pairs, but not in m9 or m10 pairs (**Figure 4 F, G, L, M, and N1-12**), and *Tb.2* and *Tb.7*, that were conversely exclusively expressed in the muscle m9 or m10 pairs (**Figure 4 D and J1-12**). Temporal variation in gene expression was also remarkable. The earliest expressed genes were *Tb.3* and *Tb.8a*, detected at the 64-cell stage (**Figure 4 E1 and K1**), immediately followed by others, such as *Tb.4b*, *Tb.8b*, and *Tb.9b* at the 100-cell stage (**Figure 4 G2, L2, and N2**). In contrast, *Tb.2*, *Tb.4a*, *Tb.5*, and *Tb.7* did not exhibit detectable expression until post-hatchling stages (**Figure 4 D9, F8, H10, and J10**). Furthermore, *Tb.5* gene showed a complex dynamic expression pattern, being strongly detected in m8 pair and weakly in the anterior cells at late hatchling stages but becoming intense in the m6 and m7 in addition of m8 by the pre-tailshift stage (**Figure 4 H10-12**). Another remarkable temporal downregulation affected *Tb.3*, *Tb.8a* and *Tb.8b* by the pre-tailshift stage, suggesting that those *Myh* genes were primarily involved in developmental processes but not in juvenile or adult muscles (**Figure 4 E12, K12, and L12**).

##### *Myh-Scp_Tb* non-muscle domains

Interestingly, some *Tb* paralogs also displayed expression in apparently non-muscle domains, suggesting possible co-option for contractile role in other tissues. *Tb.8a*, for instance, showed some clear expression domains in the trunk region during post-hatchling stages, in the areas corresponding to the developing vertical intestine and endostyle (**Figure 4 K7-11**). Additionally, at the pre-tailshift stage, the oikoblast showed expression of both Ta and several Tb paralogs (*Tb.2*, *Tb.3*, *Tb.4a*, *Tb.6*, and *Tb.8a*) (**Figure 4 B12, D12, E12, F12, I12, and K12**). Given the role of the oikoblast in house-building, this raises the possibility that these epithelial cells may possess contractile properties related to the secreting process.

## DISCUSSION

### Less muscle cells, but more tail muscle cell identities for a free-swimming lifestyle in appendicularians

The expansion and diversification of the *Myh-Scp_Tb* family reveal the presence of multiple, molecularly distinct muscle cell populations along the anteroposterior axis of the *O. dioica* tail. Each population is defined by a characteristic combinatorial expression of different *Myh-Scp_Tb* paralogues, indicating that the adult tail musculature is not a homogeneous contractile tissue, but instead comprises several muscle identities distinguished by their *Myh* repertoire. To the best of our knowledge, such a subdivision into region-specific muscle identities has not been described in ascidians, whose larval tails generate a single stereotyped oscillatory behaviour optimized for short-range dispersal before metamorphosis (Razy-Krajka & Stolfi, 2019). In contrast, *O. dioica* relies on its tail throughout its entire life cycle, using it not only for swimming but also for house inflation, house-entry and -exit, swimming, nodding, rapid escape responses, and the rhythmic water pumping through the house (Ferrández-Roldán et al., 2019; Kreneisz & Glover, 2015).

The evolution of multiple *Myh-Scp-Tb* paralogues with distinct spatial and temporal expression patterns likely enabled fine-tuned modulation of contractile properties – such as shortening velocity, force production, and fatigue resistance – within specific subsets of muscle cells. This molecular diversification provides a plausible mechanism by which a musculature composed of only ten pairs of mononucleated muscle cells can nevertheless achieve the wide repertoire of precise and energetically demanding behaviours that characterize the appendicularian pelagic lifestyle. The expression of *O. dioica*-specific duplicated paralogues of key developmental regulators, such as members of the Fgf, Wnt, and Hox families in subsets of tail muscle cells (Martí-Solans et al., 2021; Sánchez-Serna et al., 2025; Seo et al., 2004) further reinforces the view that appendicularians evolved a surprisingly complex patterning system that generates multiple muscle cell identities from a very small number of cells.

Together, these findings resolve the apparent paradox in which appendicularians, despite having far fewer tail muscle cells than ascidians, display greater behavioural sophistication. Rather than increasing cell number or relying on muscle remodelling, appendicularians appear to have evolved enhanced molecular and functional diversification within a minimal musculature, an innovation that facilitated the evolution of a fully free-swimming lifestyle.

### The loss of *Myh-Sj/bw* in appendicularians supports a pattern of regressive evolution from an ascidian-like ancestor

The loss of the *Myh-Sj/bw* type in appendicularians, which in ascidians is expressed in post-metamorphic muscles of the body wall and siphons, correlates with the absence of metamorphosis and the lack of adult-specific musculature in *O. dioica*. This correspondence suggests that post-metamorphic muscles were present in the appendicularian ancestor and were subsequently lost, together with their associated *Myh-Sj/bw* homolog, during their neotenic evolutionary transition to a fully pelagic lifestyle. In line with previous work (Ferrández-Roldán et al., 2021), these observations further support the view that the fully free-swimming appendicularian lifestyle represents a secondary derived condition, and that the last common ancestor of tunicates most likely had a biphasic ascidian-like lifestyle.

## ACKNOWLEDGEMENTS

We thank all present and past team members on CC’s laboratory for assistance and fruitful discussions, specially to Sebastian Artime Paoletti for running the Oikopleura facility in the University of Barcelona. We thank to Centres Científics i Tecnològics de la UB for sea water supply and sequencing services.

## AUTHOR CONTRIBUTIONS

Conceptualization: CC, MF; formal analysis: MF; funding acquisition: CC; investigation: MF, AF, GS, BI and CC; project administration and supervision: CC; writing: MF, CC.

## SUPPLEMENTARY FILES

**Supplementary Figure 1.**
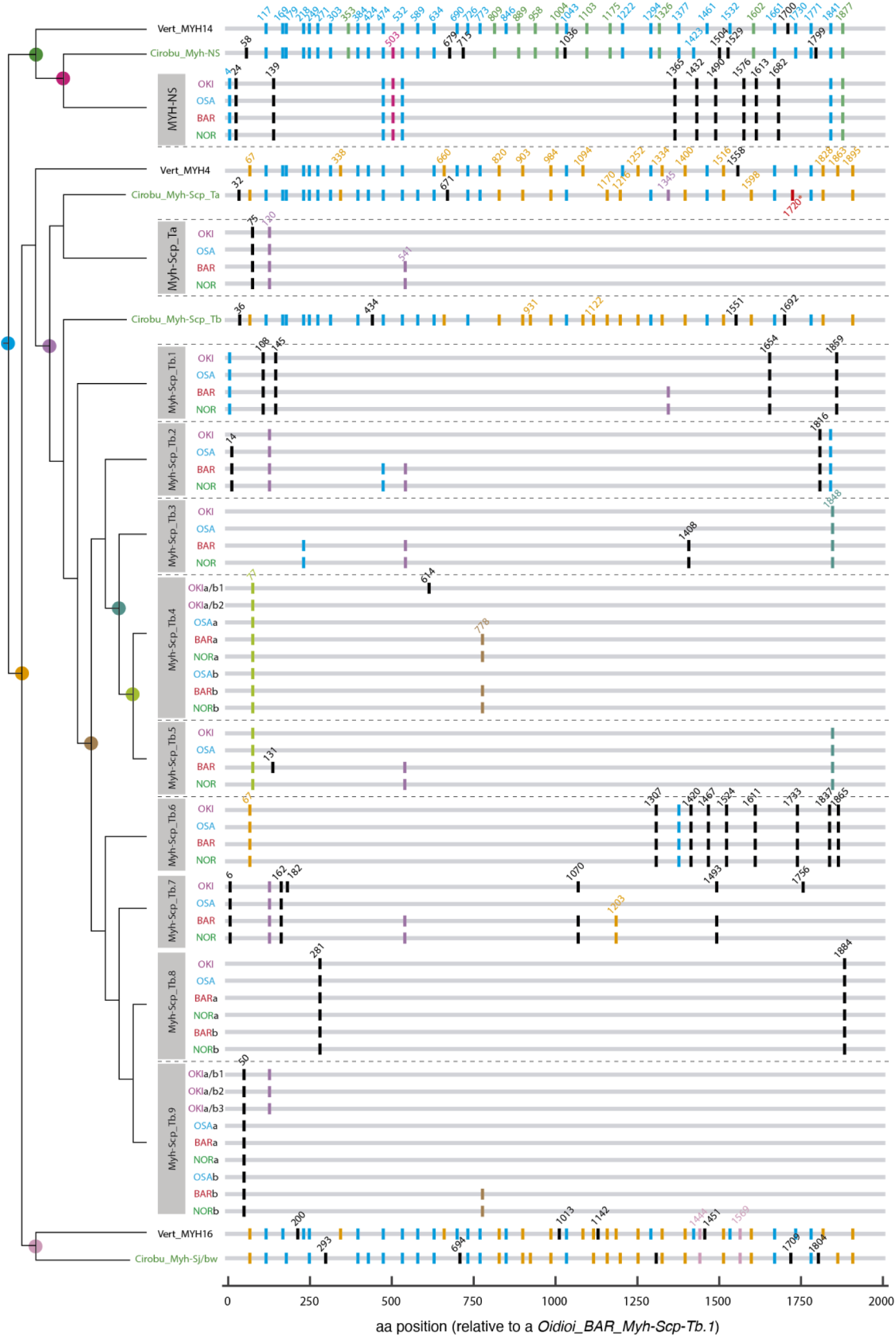
Comparative Intron analysis among *O. dioica Myh* paralog groups, and representative *Myh* genes from vertebrates and ascidians. Intron position is relative to the amino acid location in the sequence *Oidioi_BAR_Myh-Scp-Tb.1*. Introns were considered homologous when located between the same amino acids, or within a window of ±3 amino acids. Introns are color-coded according to their inferred evolutionary origin, as indicated by a coloured dot in the phylogenetic tree on the left. For example, blue indicates ancestral introns likely present before the split of *Myh-NS* and *Myh-S*, while black indicates recent, group-specific introns. The low conservation of intron organization among *O. dioica* paralog groups contrasts with the high conservation observed among ascidian and vertebrate *Myh*. Notably, all six genes identified in *C. robusta* as *Myh-Scp_Ta* showed identical organization, except for intron 170, marked in red with an asterisk.

**Supplementary Figure 2.**
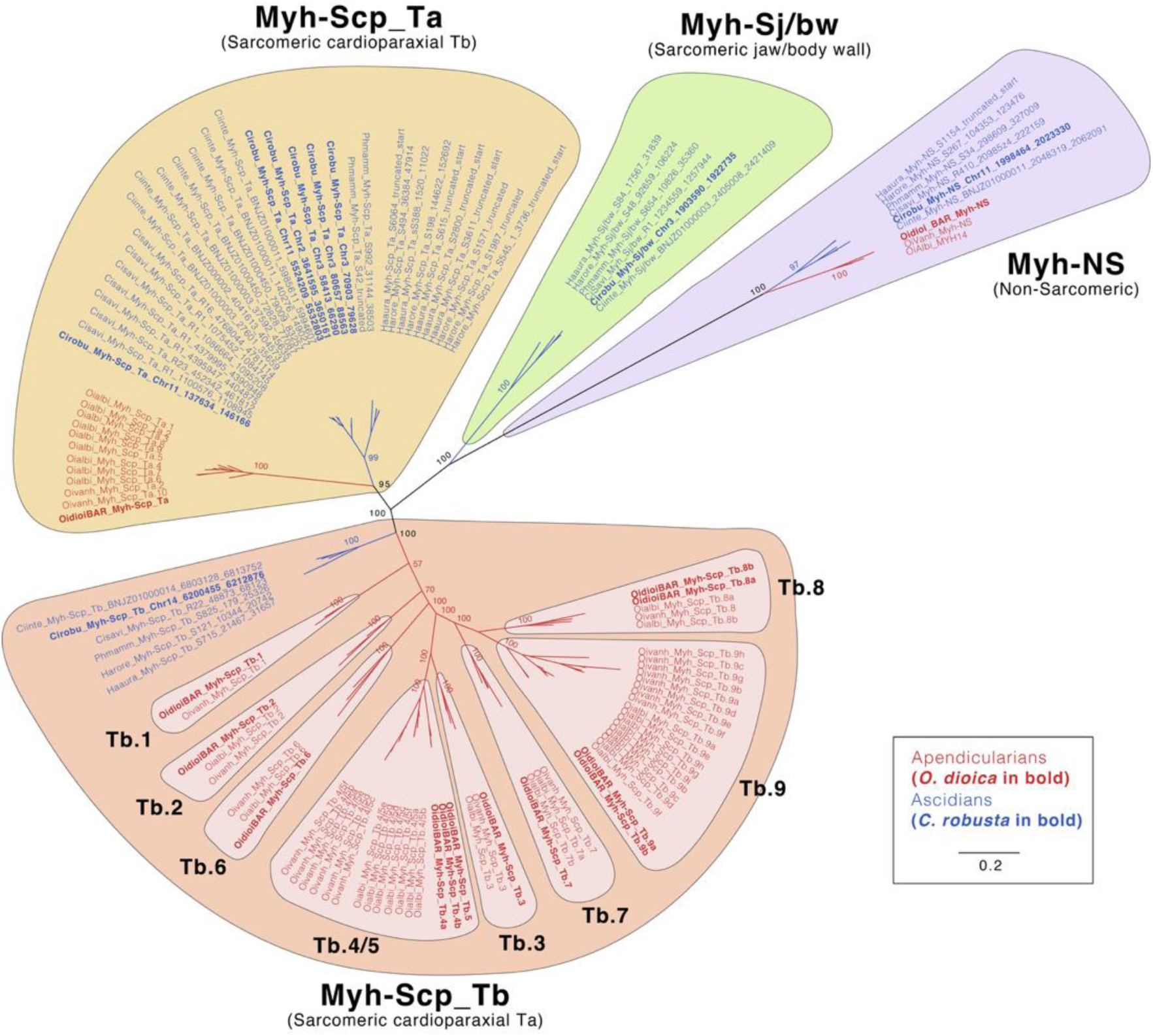
Evolutionary tree of *Myh* catalogue in tunicates. ML phylogenetic tree of the *Myh* catalogue in tunicates reveals 4 clusters, three included both ascidian (blue) and appendicularian (red) sequences, while the fourth cluster was composed exclusively of ascidian sequences. Node support values indicate ultrafast bootstrap support. The scale bar indicates amino acid substitution. Species abbreviations: Ascidian tunicates (in blue): *Ciona robusta* (Cirobu), *Ciona intestinalis* (Ciinte), *Ciona savignyi* (Cisavi), *Phallusia mammillata* (Phmamm), *Halocynthia roretzi* (Harore), *Halocynthia aurantium* (Haaura); Appendicularian tunicates (in red): *Oikopleura dioica* (Oidioi), *Oikopleura albicans* (Oialbi), *Oikopleura vanhoeffeni* (Oivanh).

**Supplementary Table 1.** *Myh* identified loci in *O. dioica* assemblies of BAR, NOR, OSA, and OKI. The high quality of the new genome assemblies enabled the identification of complete non-fragmented *Myh* loci in all cases, but one. In the OSA assembly, the gene *Myh-Scp_Tb.9a* appeared to be fragmented, with most of the sequence located on chromosome PAR (PAR:3,982,545..3,984,497(-)), while the initial portion was found on an unassembled scaffold (S187:0..2855(-)). Subsequent alignment confirmed that the ends of both fragments were completely consecutive suggesting that they belonged to the same loci, which was further corroborated by finding transcriptomic data that bridged both genomic pieces. Accordingly, we reconstructed the complete sequence by merging the two regions. For clarity, we referred to the position on chromosome PAR as the representative location of this gene in all further analyses (marked with a red asterisk), as the fragmentation likely reflects a minor miss assembly.

| Assembly | Loci | Name |
| --- | --- | --- |
| Barcelona (BAR) | XSR:10,621,546..10,633,579(-) | *Myh-NS* |
| Barcelona (BAR) | Chr2:7,396,092..7,402,033(+) | *Myh-Scp_Ta* |
| Barcelona (BAR) | PAR:5,413,769..5,420,314(-) | *Myh-Scp_Tb.1* |
| Barcelona (BAR) | Chr1:4,562,196..4,568,336(-) | *Myh-Scp_Tb.2* |
| Barcelona (BAR) | Chr2:4,993,850..5,000,646(-) | *Myh-Scp_Tb.3* |
| Barcelona (BAR) | Chr1:1,263,145..1,269,070(-) | *Myh-Scp_Tb.4a* |
| Barcelona (BAR) | Chr2:3,694,507..3,700,425(-) | *Myh-Scp_Tb.4b* |
| Barcelona (BAR) | PAR:5,687,197..5,693,214(+) | *Myh-Scp_Tb.5* |
| Barcelona (BAR) | Chr2:10,361,063..10,367,331(-) | *Myh-Scp_Tb.6* |
| Barcelona (BAR) | Chr2:273,780..280,566(+) | *Myh-Scp_Tb.7* |
| Barcelona (BAR) | XSR:7,579,141..7,585,046(+) | *Myh-Scp_Tb.8a* |
| Barcelona (BAR) | XSR:10,235,475..10,241,375(-) | *Myh-Scp_Tb.8b* |
| Barcelona (BAR) | Chr2:456,265..462,122(+) | *Myh-Scp_Tb.9a* |
| Barcelona (BAR) | PAR:12,961,103..12,967,015(-) | *Myh-Scp_Tb.9b* |
| Bergen (NOR) | scaffold_13:411,535..420,751(+) | *Myh-NS* |
| Bergen (NOR) | scaffold_5:893,345..898,044(+) | *Myh-Scp_Ta* |
| Bergen (NOR) | scaffold_64:78,898..85,330(-) | *Myh-Scp_Tb.1* |
| Bergen (NOR) | scaffold_14:57,151..62,730(-) | *Myh-Scp_Tb.2* |
| Bergen (NOR) | scaffold_159:15,552..22,487(+) | *Myh-Scp_Tb.3* |
| Bergen (NOR) | scaffold_2:1,612,242..1,619,898(-) | *Myh-Scp_Tb.4a* |
| Bergen (NOR) | scaffold_81:48,756..53,722(-) | *Myh-Scp_Tb.4b* |
| Bergen (NOR) | scaffold_19:489,542..495,995(-) | *Myh-Scp_Tb.5* |
| Bergen (NOR) | scaffold_1:1,778,608..1,785,418(-) | *Myh-Scp_Tb.6* |
| Bergen (NOR) | scaffold_60:219,316..226,145(+) | *Myh-Scp_Tb.7* |
| Bergen (NOR) | scaffold_26:142,918..148,823(-) | *Myh-Scp_Tb.8a* |
| Bergen (NOR) | scaffold_13:827,599..833,499(+) | *Myh-Scp_Tb.8b* |
| Bergen (NOR) | scaffold_7:820,732..826,638(-) | *Myh-Scp_Tb.9a* |
| Bergen (NOR) | scaffold_37:353,169..358,493(-) | *Myh-Scp_Tb.9b* |
| Osaka (OSA) | XSR:2,403,055..2,414,830(+) | *Myh-NS* |
| Osaka (OSA) | Chr2:7,509,565..7,503,659(+) | *Myh-Scp_Ta* |
| Osaka (OSA) | Chr2:11,751,140..11,757,586(+) | *Myh-Scp_Tb.1* |
| Osaka (OSA) | Chr1:9,473,201..9,479,204(+) | *Myh-Scp_Tb.2* |
| Osaka (OSA) | Chr2:12,1010,871..12,107,785(+) | *Myh-Scp_Tb.3* |
| Osaka (OSA) | PAR:1,628,384..1,634,259(-) | *Myh-Scp_Tb.4a* |
| Osaka (OSA) | PAR:4,680,989..4,686,874(+) | *Myh-Scp_Tb.4b* |
| Osaka (OSA) | PAR:14,913,622..14,919,558(-) | *Myh-Scp_Tb.5* |
| Osaka (OSA) | Chr2:12,990,815..12,997,272(-) | *Myh-Scp_Tb.6* |
| Osaka (OSA) | Chr2:4,780,330..4,786,985(+) | *Myh-Scp_Tb.7* |
| Osaka (OSA) | PAR:12,872,721..12,878,618(+) | *Myh-Scp_Tb.8* |
| Osaka (OSA) | *PAR:3,982,545..3,984,497(-) | *Myh-Scp_Tb.9a* |
| Osaka (OSA) | PAR:15,242,514..15,248,369(-) | *Myh-Scp_Tb.9b* |
| Okinawa (OKI) | XSR:3,684,116..3,692,291(+) | *Myh-NS* |
| Okinawa (OKI) | Chr2:6,478,579..6,472,666(+) | *Myh-Scp_Ta* |
| Okinawa (OKI) | Chr2:9,186,247..9,192,257(+) | *Myh-Scp_Tb.1* |
| Okinawa (OKI) | PAR:2,764,739..2,770,674(+) | *Myh-Scp_Tb.2* |
| Okinawa (OKI) | Chr2:9,688,868..9,694,749(+) | *Myh-Scp_Tb.3* |
| Okinawa (OKI) | Chr2:11,893,510..11,899,636(-) | *Myh-Scp_Tb.4a/b1* |
| Okinawa (OKI) | Chr2:1,391,391..1,397,274(-) | *Myh-Scp_Tb.4a/b2* |
| Okinawa (OKI) | Chr2:12,694,958..12,700,881(-) | *Myh-Scp_Tb.5* |
| Okinawa (OKI) | Chr2:9,222,112..9,228,400(-) | *Myh-Scp_Tb.6* |
| Okinawa (OKI) | Chr2:2,811,282..2,820,152(+) | *Myh-Scp_Tb.7* |
| Okinawa (OKI) | XSR:580,993..586,863(+) | *Myh-Scp_Tb.8* |
| Okinawa (OKI) | Chr2:8,034,804..8,040,696(+) | *Myh-Scp_Tb.9a/b1* |
| Okinawa (OKI) | PAR:14,030,606..14,036,503(-) | *Myh-Scp_Tb.9a/b2* |
| Okinawa (OKI) | Chr1:11,825,587..11,831,481(-) | *Myh-Scp_Tb.9a/b3* |

**Supplementary Table 2.**
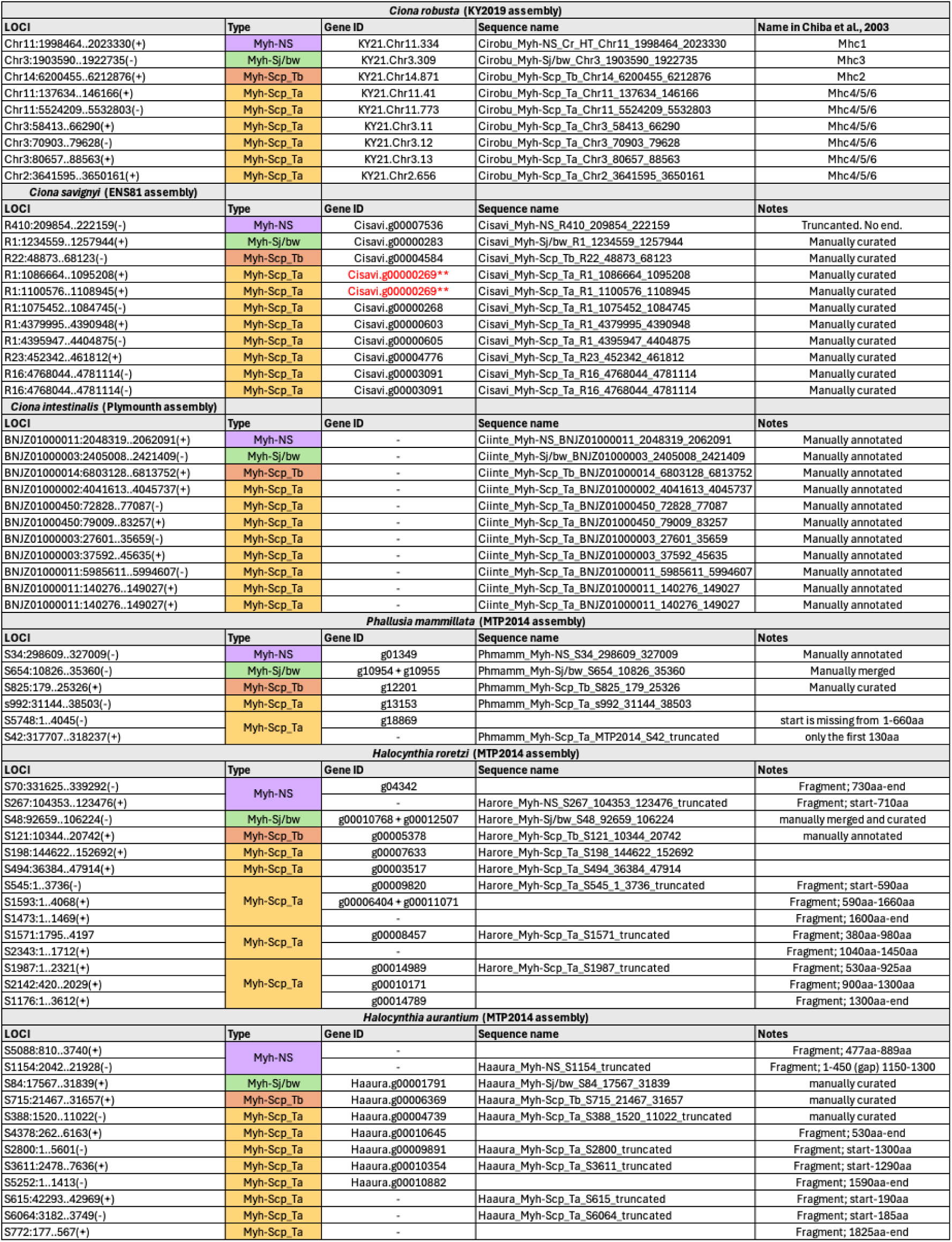
List of loci identified in ascidian species (*Ciona robusta, Ciona intestinalis, Ciona savignyi, Phallusia mammillata, Halocynthia roretzi, Halocynthia aurantium*). Only loci containing a conserved region of the Myh head domain were considered for phylogenetic analysis. For fragmented loci, the position of the fragment relative to the overall alignment of all proteins is indicated, considering a total protein length of approximately 2000 amino acids.

**Supplementary Table 3.** List of loci identified in appendicularian species (*Oikopleura albicans and Oikopleura vanhoeffeni*). Only loci containing a conserved region of the Myh head domain were considered for phylogenetic analysis. Discarded loci have been indicated by a dash (-) in the sequence name column.

| *Oikopleura vanhoeffeni* |  |  |  |
| --- | --- | --- | --- |
| LOCI | Type | Sequence name | Notes |
| SCLH01001028:175KB..185KB(+) | Myh-NS | Oivanh_Myh-NS | start |
| SCLH01001615:48KB..60KB(-) |  | - | end |
| SCLH01000560:234KB..249KB(+) | Myh-Scp_Tb | Oivanh_Myh-Scp_Tb.8a | manually annotated |
| SCLH01000395:163KB..173KB(+) | Myh-Scp_Tb | Oivanh_Myh-Scp_Tb.9a | manually annotated |
| SCLH01000581:422KB..431KB(-) | Myh-Scp_Tb | Oivanh_Myh-Scp_Tb.9b | manually annotated |
| SCLH01000581:398KB..407KB(-) | Myh-Scp_Tb | Oivanh_Myh-Scp_Tb.9c | manually annotated |
| SCLH01000943:231KB..241KB(-) | Myh-Scp_Tb | Oivanh_Myh-Scp_Tb.9d | manually annotated |
| SCLH01000153:170KB..180KB(-) | Myh-Scp_Tb | Oivanh_Myh-Scp_Tb.9e | manually annotated |
| SCLH01001199:62KB..70KB(-) | Myh-Scp_Tb | Oivanh_Myh-Scp_Tb.9f | manually annotated |
| SCLH01000581:411KB..417KB(-) | Myh-Scp_Tb | Oivanh_Myh-Scp_Tb.9g | Fragment; start-1380aa |
| SCLH01001158:12KB..44KB(-) | Myh-Scp_Tb | Oivanh_Myh-Scp_Tb.9h | Fragment; start-1115aa |
| SCLH01003921:3KB..5KB(-) | Myh-Scp_Tb | - | Fragment; 1660aa-end |
| SCLH01000404:177KB..210KB(-) | Myh-Scp_Tb | Oivanh_Myh-Scp_Tb.7a | manually annotated |
| SCLH01001210:88KB..89KB(+) | Myh-Scp_Tb | - | Fragment; 390aa-550aa |
| SCLH01002236:22KB..60KB(+) | Myh-Scp_Tb | Oivanh_Myh-Scp_Tb.2 | manually annotated |
| SCLH01000012:65KB..82KB(-) | Myh-Scp_Tb | Oivanh_Myh-Scp_Tb.3 | manually annotated |
| SCLH01000369:393KB..405KB(+) | Myh-Scp_Tb | Oivanh_Myh-Scp_Tb.4/5a | manually annotated |
| SCLH01000369:352KB..367KB(-) | Myh-Scp_Tb | Oivanh_Myh-Scp_Tb.4/5b | manually annotated |
| SCLH01000943:324KB..334KB(+) | Myh-Scp_Tb | Oivanh_Myh-Scp_Tb.4/5c | manually annotated |
| SCLH01000118:146KB..156KB(+) | Myh-Scp_Tb | Oivanh_Myh-Scp_Tb.4/5d | manually annotated |
| SCLH01000391:135KB..140KB(-) | Myh-Scp_Tb | Oivanh_Myh-Scp_Tb.4/5e | Fragment; start-500aa |
| SCLH01063333:25KB..30KB(-) | Myh-Scp_Tb | - | Fragment; 505aa-end |
| SCLH01000448:481KB..486KB(+) | Myh-Scp_Tb | Oivanh_Myh-Scp_Tb.4/5f | Fragment; start-355aa |
| SCLH01000288:402KB..407KB(-) | Myh-Scp_Tb | - | Fragment; 350aa-765aa |
| SCLH01048532:0KB..1KB(-) | Myh-Scp_Tb | - | Fragment; 1480aa-1610aa |
| SCLH01000212:54KB..63KB(+) | Myh-Scp_Tb | Oivanh_Myh-Scp_Tb.6 | Fragment; start-500aa |
| SCLH01049440:1KB..2KB(-) | Myh-Scp_Tb | - | Fragment; 500aa-610aa |
| SCLH01062268:0KB..1KB(+) | Myh-Scp_Tb | - | Fragment; 605aa-725aa |
| SCLH01002204:151KB..185KB(-) | Myh-Scp_Tb | Oivanh_Myh-Scp_Tb.1 | manually annotated |
| SCLH01000378:35KB..45KB(+) | Myh-Scp_Ta | Oivanh_Myh-Scp_Ta.1 | manually annotated |
| SCLH01000403:51KB..62KB(-) | Myh-Scp_Ta | Oivanh_Myh-Scp_Ta.2 | manually annotated |
| *Oikopleura albicans* |  |  |  |
| LOCI | Type | Sequence name | Notes |
| SCLG01000124:36KB..50KB | Myh-NS | Oialbi_Myh-NS | manually annotated |
| SCLG01000056:71KB..79KB | Myh-Scp_Tb | Oialbi_Myh-Scp_Tb.8a | manually annotated |
| SCLG01000062:354KB..366KB | Myh-Scp_Tb | Oialbi_Myh-Scp_Tb.8b | manually annotated |
| SCLG01000142:304KB..314KB | Myh-Scp_Tb | Oialbi_Myh-Scp_Tb.9a | manually annotated |
| SCLG01000022:261KB..268KB | Myh-Scp_Tb | Oialbi_Myh-Scp_Tb.9b | manually annotated |
| SCLG01000050:1327KB..1334KB | Myh-Scp_Tb | Oialbi_Myh-Scp_Tb.9c | manually annotated |
| SCLG01000283:155KB..163KB | Myh-Scp_Tb | Oialbi_Myh-Scp_Tb.9d | manually annotated |
| SCLG01000049:553KB..560KB | Myh-Scp_Tb | Oialbi_Myh-Scp_Tb.9e | manually annotated |
| SCLG01000595:57KB..64KB | Myh-Scp_Tb | Oialbi_Myh-Scp_Tb.9f | manually annotated |
| SCLG01013153:4KB..8KB | Myh-Scp_Tb | Oialbi_Myh-Scp_Tb.9g | Fragment; start-450aa |
| SCLG01014172:0KB..3KB | Myh-Scp_Tb | Oialbi_Myh-Scp_Tb.9h | Fragment; 220aa-665aa |
| SCLG01000595:90KB..92KB | Myh-Scp_Tb | - | Fragment; start-135aa |
| SCLG01014433:2KB..5KB | Myh-Scp_Tb | Oialbi_Myh-Scp_Tb.9i | Fragment; 220aa-490aa |
| SCLG01018614:0KB..2KB | Myh-Scp_Tb | - | Fragment; 590aa-970aa |
| SCLG01008761:4KB..7KB | Myh-Scp_Tb | - | manually annotated |
| SCLG01000086:291KB..305KB | Myh-Scp_Tb | Oialbi_Myh-Scp_Tb.7a | manually annotated |
| SCLG01009445:2KB..5KB | Myh-Scp_Tb | Oialbi_Myh-Scp_Tb.7b | Fragment; 360aa-830aa |
| SCLG01016459:0KB..2KB | Myh-Scp_Tb | - | Fragment; 960aa-1370aa |
| SCLG01000623:25KB..33KB | Myh-Scp_Tb | Oialbi_Myh-Scp_Tb.2 | manually annotated |
| SCLG01000014:290KB..299KB | Myh-Scp_Tb | Oialbi_Myh-Scp_Tb.3 | manually annotated |
| SCLG01000833:18KB..27KB | Myh-Scp_Tb | Oialbi_Myh-Scp_Tb.4/5a | manually annotated |
| SCLG01000142:92KB..99KB | Myh-Scp_Tb | Oialbi_Myh-Scp_Tb.4/5b | manually annotated |
| SCLG01000123:100KB..108KB | Myh-Scp_Tb | Oialbi_Myh-Scp_Tb.4/5c | manually annotated |
| SCLG01000178:512KB..521KB | Myh-Scp_Tb | Oialbi_Myh-Scp_Tb.4/5d | manually annotated |
| SCLG01000003:88KB..95KB | Myh-Scp_Tb | Oialbi_Myh-Scp_Tb.4/5e | manually annotated |
| SCLG01000049:541KB..548KB | Myh-Scp_Tb | Oialbi_Myh-Scp_Tb.4/5f | manually annotated |
| SCLG01002169:0KB..5KB | Myh-Scp_Tb | Oialbi_Myh-Scp_Tb.4/5g | manually annotated |
| SCLG01030508:0KB..2KB | Myh-Scp_Tb | - | Fragment; 115aa-300aa |
| SCLG01000091:41KB..59KB | Myh-Scp_Tb | Oialbi_Myh-Scp_Tb.6 | manually annotated |
| SCLG01000091:40KB..52KB | Myh-Scp_Tb | - | Fragment; start-860aa |
| SCLG01019803:0KB..4KB | Myh-Scp_Tb | - | Fragment; start-860aa |
| SCLG01000292:117KB..125KB | Myh-Scp_Ta | Oialbi_Myh-Scp_Ta.1 | manually annotated |
| SCLG01000005:742KB..749KB | Myh-Scp_Ta | Oialbi_Myh-Scp_Ta.2 | manually annotated |
| SCLG01000081:79KB..90KB | Myh-Scp_Ta | Oialbi_Myh-Scp_Ta.3 | manually annotated |
| SCLG0100169:534KB..542KB | Myh-Scp_Ta | Oialbi_Myh-Scp_Ta.4 | manually annotated |
| SCLG01000030:7KB..15KB | Myh-Scp_Ta | Oialbi_Myh-Scp_Ta.5 | manually annotated |
| SCLG01000221:203KB..213KB | Myh-Scp_Ta | Oialbi_Myh-Scp_Ta.6 | manually annotated |
| SCLG01030282:0KB..1KB | Myh-Scp_Ta | Oialbi_Myh-Scp_Ta.7 | Fragment; 560aa-875aa |
| SCLG01015296:0KB..2KB | Myh-Scp_Ta | Oialbi_Myh-Scp_Ta.8 | Fragment; 415aa-930aa |
| SCLG01030447:0KB..2KB | Myh-Scp_Ta | Oialbi_Myh-Scp_Ta.9 | Fragment; 570aa-928aa |
| SCLG01031245:0KB..1KB | Myh-Scp_Ta | Oialbi_Myh-Scp_Ta.10 | Fragment; 480aa-790aa |
| SCLG01000194:1KB..3KB | Myh-Scp_Ta | - | Fragment; 995aa-1580aa |
| SCLG01005058:2K..4K | Myh-Scp_Ta | - | Fragment; 1520aa-1925aa |
| SCLG01005058:2K..4K | Myh-Scp_Ta | - | Fragment; 1520aa-1925aa |

**Supplementary Table 4.**
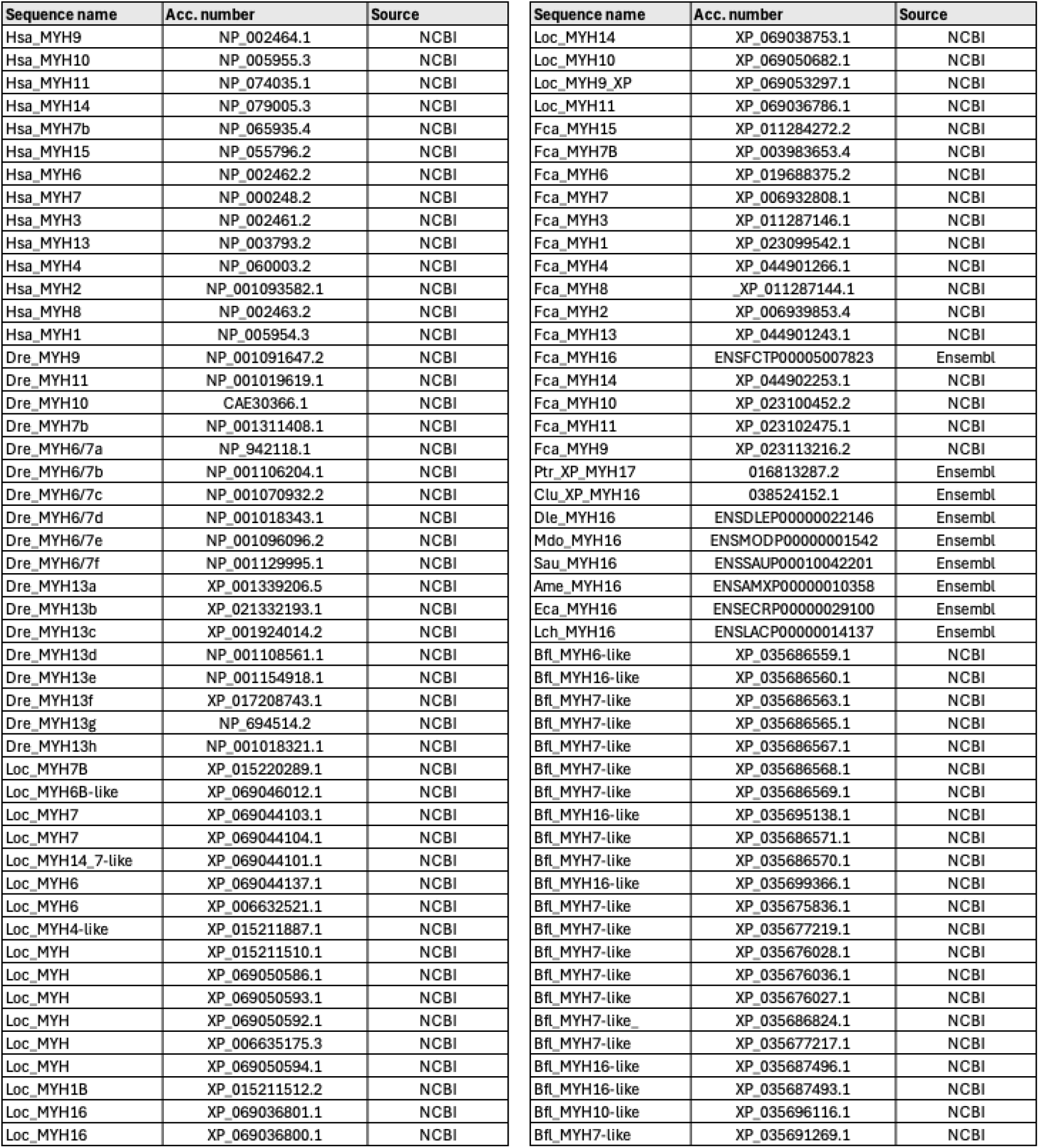
List of vertebrates and cephalochordates sequences used for phylogenetic analysis of the Myh gene evolution within chordates.

**Supplementary Table 5.**
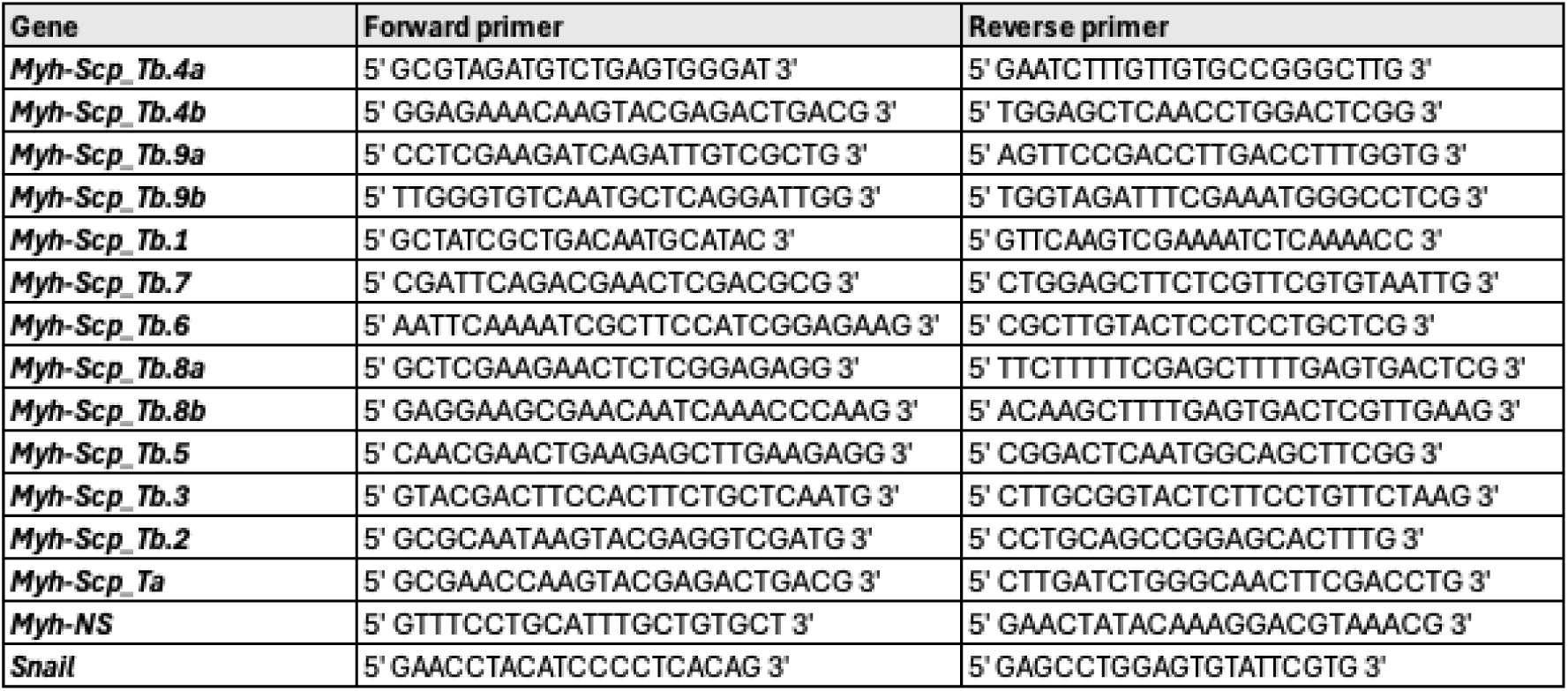
List of the primers designed for the cloning of *Myh* gene catalogue.

## BIBLIOGRAPHY

Albalat, R., & Cañestro, C. (2016). Evolution by gene loss. In Nature Reviews Genetics (Vol. 17, Issue 7, pp. 379–391). Nature Publishing Group. 10.1038/nrg.2016.39

Bassham, S., & Postlethwait, J. (2000). Brachyury (T) expression in embryos of a larvacean urochordate, Oikopleura dioica, and the ancestral role of T. Dev Biol, 220(2)), 322-32. 10.1006/dbio.2000.9647

Bone, Q. (Ed.). (1998). The Biology of Pelagic Tunicates. Oxford University Press Oxford. 10.1093/oso/9780198540243.001.0001

Bourlat, S. J., Juliusdottir, T., Lowe, C. J., Freeman, R., Aronowicz, J., Kirschner, M., Lander, E. S., Thorndyke, M., Nakano, H., Kohn, A. B., Heyland, A., Moroz, L. L., Copley, R. R., & Telford, M. J. (2006). Deuterostome phylogeny reveals monophyletic chordates and the new phylum Xenoturbellida. Nature, 444(7115), 85–88. 10.1038/nature05241

Brozovic, M., Dantec, C., Dardaillon, J., Dauga, D., Faure, E., Gineste, M., Louis, A., Naville, M., Nitta, K. R., Piette, J., Reeves, W., Scornavacca, C., Simion, P., Vincentelli, R., Bellec, M., Aicha, S. Ben, Fagotto, M., Guéroult-Bellone, M., Haeussler, M., … Lemaire, P. (2018). ANISEED 2017: Extending the integrated ascidian database to the exploration and evolutionary comparison of genome-scale datasets. Nucleic Acids Research, 46(D1), D718–D725. 10.1093/nar/gkx1108

Cañestro, C., Yokoi, H., & Postlethwait, J. H. (2007). Evolutionary developmental biology and genomics. Nature Reviews. Genetics, 8(12), 932–942. 10.1038/nrg2226

Chiba, S., Awazu, S., Itoh, M., Chin-Bow, S. T., Satoh, N., Satou, Y., & Hastings, K. E. M. (2003). A genomewide survey of developmentally relevant genes in Ciona intestinalis. Development Genes and Evolution, 213(5–6), 291–302. 10.1007/s00427-003-0324-x

Danks, G., Campsteijn, C., Parida, M., Butcher, S., Doddapaneni, H., Fu, B., Petrin, R., Metpally, R., Lenhard, B., Wincker, P., Chourrout, D., Thompson, E. M., & Manak, J. R. (2013). OikoBase: A genomics and developmental transcriptomics resource for the urochordate Oikopleura dioica. Nucleic Acids Research, 41(D1), 1–9. 10.1093/nar/gks1159

Delsuc, F., Brinkmann, H., Chourrout, D., & Philippe, H. (2006). Tunicates and not cephalochordates are the closest living relatives of vertebrates. Nature, 439(7079), 965–968. 10.1038/nature04336

Delsuc, F., Philippe, H., Tsagkogeorga, G., Simion, P., Tilak, M. K., Turon, X., López-Legentil, S., Piette, J., Lemaire, P., & Douzery, E. J. P. (2018). A phylogenomic framework and timescale for comparative studies of tunicates. BMC Biology, 16(1). 10.1186/s12915-018-0499-2

Denoeud, F., Henriet, S., Mungpakdee, S., Aury, J.-M., Da Silva, C., Brinkmann, H., Mikhaleva, J., Olsen, L. C., Jubin, C., Cañestro, C., Bouquet, J.-M., Danks, G., Poulain, J., Campsteijn, C., Adamski, M., Cross, I., Yadetie, F., Muffato, M., Louis, A., … Chourrout, D. (2010). Plasticity of Animal Genome Architecture Unmasked by Rapid Evolution of a Pelagic Tunicate. Science, 330(6009), 1381–1385. 10.1126/science.1194167

Fenaux, R. (1986). The house of Oikopleura dioica (Tunicata, Appendicularia): Structure and functions. Zoomorphology, 106(4), 224–231. 10.1007/bf00312043

Fenaux, R. (1998). Anatomy and functional morphology of the Appendicularia. In Q. Bone (Ed.), The Biology of Pelagic Tunicates (pp. 25–34). Oxford University Press.

Ferrández-Roldán, A., Fabregà-Torrus, M., Sánchez-Serna, G., Duran-Bello, E., Joaquín-Lluís, M., Bujosa, P., Plana-Carmona, M., Garcia-Fernàndez, J., Albalat, R., & Cañestro, C. (2021). Cardiopharyngeal deconstruction and ancestral tunicate sessility. Nature, 599(7885), 431–435. 10.1038/s41586-021-04041-w

Ferrández-Roldán, A., Martí-Solans, J., Cañestro, C., & Albalat, R. (2019). Oikopleura dioica: An Emergent Chordate Model to Study the Impact of Gene Loss on the Evolution of the Mechanisms of Development. In W. Tworzydlo & S. Bilinski (Eds.), Evo-Devo: Non-model Species in Cell and Developmental Biology. (Vol. 68, pp. 63–105). Springer, Cham. 10.1007/978-3-030-23459-1_4

Flood, P. R., & Deibel, D. (1998). The appendicularian house. In Q. Bone (Ed.), The Biology of Pelagic Tunicates (pp. 105–137). Oxford University Press.

Fuchs, B., Wang, W., Graspeuntner, S., Li, Y., Insua, S., Herbst, E.-M., Dirksen, P., Böhm, A.-M., Hemmrich, G., Sommer, F., Domazet-Lošo, T., Klostermeier, U. C., Anton-Erxleben, F., Rosenstiel, P., Bosch, T. C. G., & Khalturin, K. (2014). Regulation of polyp-to-jellyfish transition in Aurelia aurita. Current Biology : CB, 24(3), 263–273. 10.1016/j.cub.2013.12.003

Guindon, S., Dufayard, J.-F., Lefort, V., Anisimova, M., Hordijk, W., & Gascuel, O. (2010). New algorithms and methods to estimate maximum-likelihood phylogenies: assessing the performance of PhyML 3.0. Systematic Biology, 59(3), 307–321. 10.1093/sysbio/syq010

Johnson, C. J., Razy-Krajka, F., & Stolfi, A. (2020). Expression of smooth muscle-like effectors and core cardiomyocyte regulators in the contractile papillae of Ciona. EvoDevo, 11(1), 15. 10.1186/s13227-020-00162-x

Kalyaanamoorthy, S., Minh, B. Q., Wong, T. K. F., von Haeseler, A., & Jermiin, L. S. (2017). ModelFinder: fast model selection for accurate phylogenetic estimates. Nature Methods, 14(6), 587–589. 10.1038/nmeth.4285

Kocot, K. M., Tassia, M. G., Halanych, K. M., & Swalla, B. J. (2018). Phylogenomics offers resolution of major tunicate relationships. Molecular Phylogenetics and Evolution, 121, 166–173. 10.1016/j.ympev.2018.01.005

Kreneisz, O., & Glover, J. C. (2015). Developmental Characterization of Tail Movements in the Appendicularian Urochordate Oikopleura dioica. Brain, Behavior and Evolution, 86(3–4), 191–209. 10.1159/000439517

Larsson, A. (2014). AliView: A fast and lightweight alignment viewer and editor for large datasets. Bioinformatics, 30(22), 3276–3278. 10.1093/bioinformatics/btu531

Laudet, V. (2011). The origins and evolution of vertebrate metamorphosis. Current Biology : CB, 21(18), R726–37. 10.1016/j.cub.2011.07.030

Lemaire, P. (2011). Evolutionary crossroads in developmental biology: The tunicates. Development, 138(11), 2143–2152. 10.1242/dev.048975

Lemoine, F., Correia, D., Lefort, V., Doppelt-Azeroual, O., Mareuil, F., Cohen-Boulakia, S., & Gascuel, O. (2019). NGPhylogeny.fr: new generation phylogenetic services for non-specialists. Nucleic Acids Research, 47(W1), W260–W265. 10.1093/nar/gkz303

Martí-Solans, J., Belyaeva, O. V., Torres-Aguila, N. P., Kedishvili, N. Y., Albalat, R., & Cañestro, C. (2016). Coelimination and Survival in Gene Network Evolution: Dismantling the RA-Signaling in a Chordate. Molecular Biology and Evolution, 33(9), 2401–2416. 10.1093/molbev/msw118

Martí-Solans, J., Ferrández-Roldán, A., Godoy-Marín, H., Badia-Ramentol, J., Torres-Aguila, N. P., Rodríguez-Marí, A., Bouquet, J. M., Chourrout, D., Thompson, E. M., Albalat, R., & Cañestro, C. (2015). Oikopleura dioica culturing made easy: a low-cost facility for an emerging animal model in EvoDevo. Genesis (New York, N.Y. : 2000), 53(1), 183–193. 10.1002/dvg.22800

Martí-Solans, J., Godoy-Marín, H., Diaz-Gracia, M., Onuma, T. A., Nishida, H., Albalat, R., & Cañestro, C. (2021). Massive Gene Loss and Function Shuffling in Appendicularians Stretch the Boundaries of Chordate Wnt Family Evolution. Frontiers in Cell and Developmental Biology, 9, 700827. 10.3389/fcell.2021.700827

Masunaga, A., Mansfield, M. J., Tan, Y., Liu, A. W., Bliznina, A., Barzaghi, P., Hodgetts, T. L., Ferrández-Roldán, A., Cañestro, C., Onuma, T. A., Plessy, C., & Luscombe, N. M. (2022). The cosmopolitan appendicularian Oikopleura dioica reveals hidden genetic diversity around the globe. Marine Biology, 169(12), 157. 10.1007/s00227-022-04145-5

McGuigan, K. (2004). Evolution of Sarcomeric Myosin Heavy Chain Genes: Evidence from Fish. Molecular Biology and Evolution, 21(6), 1042–1056. 10.1093/molbev/msh103

Mikhaleva, Y., Skinnes, R., Sumic, S., Thompson, E. M., & Chourrout, D. (2018). Development of the house secreting epithelium, a major innovation of tunicate larvaceans, involves multiple homeodomain transcription factors. Developmental Biology, 443(2), 117–126. 10.1016/j.ydbio.2018.09.006

Nakayama-Ishimura, A., Chambon, J., Horie, T., Satoh, N., & Sasakura, Y. (2009). Delineating metamorphic pathways in the ascidian Ciona intestinalis. Developmental Biology, 326(2), 357–367. 10.1016/j.ydbio.2008.11.026

Naville, M., Henriet, S., Warren, I., Sumic, S., Reeve, M., Volff, J. N., & Chourrout, D. (2019). Massive Changes of Genome Size Driven by Expansions of Non-autonomous Transposable Elements. Current Biology, 29(7), 1161–1168. 10.1016/j.cub.2019.01.080

Nguyen, L.-T., Schmidt, H. A., von Haeseler, A., & Minh, B. Q. (2015). IQ-TREE: A Fast and Effective Stochastic Algorithm for Estimating Maximum-Likelihood Phylogenies. Molecular Biology and Evolution, 32(1), 268–274. 10.1093/molbev/msu300

Plessy, C., Mansfield, M. J., Bliznina, A., Masunaga, A., West, C., Tan, Y., Liu, A. W., Grašič, J., del Río Pisula, M. S., Sánchez-Serna, G., Fabrega-Torrus, M., Ferrández-Roldán, A., Roncalli, V., Navratilova, P., Thompson, E. M., Onuma, T., Nishida, H., Cañestro, C., & Luscombe, N. M. (2024). Extreme genome scrambling in marine planktonic *Oikopleura dioica* cryptic species. Genome Research. 10.1101/gr.278295.123

Prünster, M. M., Ricci, L., Brown, F. D., & Tiozzo, S. (2019). Modular co-option of cardiopharyngeal genes during non-embryonic myogenesis. EvoDevo, 10(1), 3. 10.1186/s13227-019-0116-7

Razy-Krajka, F., & Stolfi, A. (2019). Regulation and evolution of muscle development in tunicates. EvoDevo, 10(1), 1–34. 10.1186/s13227-019-0125-6

Sánchez-Serna, G., Badia-Ramentol, J., Bujosa, P., Ferrández-Roldán, A., Torres-Águila, N. P., Fabregà-Torrus, M., Wibisana, J. N., Mansfield, M. J., Plessy, C., Luscombe, N. M., Albalat, R., & Cañestro, C. (2025). Less, but More: New Insights From Appendicularians on Chordate Fgf Evolution and the Divergence of Tunicate Lifestyles. Molecular Biology and Evolution, 42(1). 10.1093/molbev/msae260

Satoh, N. (1994). Developmental Biology of Ascidians. Cambridge University Press.

Schiaffino, S., & Reggiani, C. (2011). Fiber Types in Mammalian Skeletal Muscles. Physiological Reviews, 91(4), 1447–1531. 10.1152/physrev.00031.2010

Seo, H.-C., Edvardsen, R. B., Maeland, A. D., Bjordal, M., Jensen, M. F., Hansen, A., Flaat, M., Weissenbach, J., Lehrach, H., Wincker, P., Reinhardt, R., & Chourrout, D. (2004). Hox cluster disintegration with persistent anteroposterior order of expression in Oikopleura dioica. Nature, 431(7004), 67–71. 10.1038/nature02709

Singh, T. R., Tsagkogeorga, G., Delsuc, F., Blanquart, S., Shenkar, N., Loya, Y., Douzery, E. J., & Huchon, D. (2009). Tunicate mitogenomics and phylogenetics: peculiarities of the Herdmania momus mitochondrial genome and support for the new chordate phylogeny. BMC Genomics, 10(1), 534. 10.1186/1471-2164-10-534

Stach, T. (2002). Phylogeny of Tunicata inferred from molecular and morphological characters. Molecular Phylogenetics and Evolution, 25(3), 408–428. 10.1016/S1055-7903(02)00305-6

Stach, T., Winter, J., Bouquet, J.-M., Chourrout, D., & Schnabel, R. (2008). Embryology of a planktonic tunicate reveals traces of sessility. www.pnas.org/cgi/content/full/

Stedman, H. H., Kozyak, B. W., Nelson, A., Thesier, D. M., Su, L. T., Low, D. W., Bridges, C. R., Shrager, J. B., Minugh-Purvis, N., & Mitchell, M. A. (2004). Myosin gene mutation correlates with anatomical changes in the human lineage. www.nature.com/nature

Swalla, B. J., Cameron, C. B., Corley, L. S., & Garey, J. R. (2000). Urochordates Are Monophyletic Within the Deuterostomes. Systematic Biology, 49(1), 52–64. 10.1080/10635150050207384

Szathmáry, E. (2015). Toward major evolutionary transitions theory 2.0. Proceedings of the National Academy of Sciences of the United States of America, 112(33), 10104–10111. 10.1073/pnas.1421398112

Wada, H. (1998). Evolutionary History of Free-Swimming and Sessile Lifestyles in Urochordates as Deduced from 18S rDNA Molecular Phylogeny. Mol. Biol. Evol, 15(9), 1189–1194. https://academic.oup.com/mbe/article/15/9/1189/1412733

Weiss, A., McDonough, D., Wertman, B., Acakpo-Satchivi, L., Montgomery, K., Kucherlapati, R., Leinwand, L., & Krauter, K. (1999). Organization of human and mouse skeletal myosin heavy chain gene clusters is highly conserved. Proceedings of the National Academy of Sciences, 96(6), 2958–2963. 10.1073/pnas.96.6.2958

Yokobori, S., Kurabayashi, A., Neilan, B. A., Maruyama, T., & Hirose, E. (2006). Multiple origins of the ascidian-Prochloron symbiosis: Molecular phylogeny of photosymbiotic and non-symbiotic colonial ascidians inferred from 18S rDNA sequences. Molecular Phylogenetics and Evolution, 40(1), 8–19. 10.1016/j.ympev.2005.11.025

